# Plasticity of the electrical connectome of *C. elegans*

**DOI:** 10.1101/406207

**Authors:** Abhishek Bhattacharya, Ulkar Aghayeva, Emily Berghoff, Oliver Hobert

## Abstract

The patterns of electrical synapses of an animal nervous system (“electrical connectome”), as well as the functional properties and plasticity of electrical synapses, are defined by the neuron type-specific complement of electrical synapse constituents. We systematically examine here properties of the electrical connectome of the nematode *C. elegans* through a genome- and nervous system-wide analysis of the expression pattern of the central components of invertebrate electrical synapses, the innexins, revealing highly complex combinatorial patterns of innexin expression throughout the nervous system. We find that the complex expression patterns of 12 out of 14 neuronally expressed innexins change in a strikingly neuron type-specific manner throughout most of the nervous system, if animals encounter harsh environmental conditions and enter the dauer arrest stage. We systematically describe the plasticity of locomotory patterns of dauer stage animals and, by analyzing several individual electrical synapses, we demonstrate that dauer stage-specific electrical synapse remodeling is responsible for specific aspects of the altered locomotory patterns as well as altered chemosensory behavior of dauer stage animals. We describe an intersectional gene regulatory mechanism, involving terminal selector and FoxO transcription factors that are responsible for inducing innexin expression changes in a neuron type- and environment-specific manner. Taken together, our studies illustrate the remarkably dynamic nature of electrical synapses on a nervous system-wide level and describe regulatory strategies for how these alterations are achieved.

## INTRODUCTION

Understanding the detailed anatomical synaptic wiring of an entire nervous system - its ‘connectome’ - is an essential first step to elucidate how a nervous system processes information and generates behavior. The adult *C. elegans* hermaphrodite was the first organism for which an entire connectome has been established (White et al., 1986), recently followed by that of a simple chordate, *Ciona intestinalis* (Ryan et al., 2016), and similar efforts are now underway for the *Drosophila* and mouse brains (Helmstaedter et al., 2013; Kasthuri et al., 2015; Takemura et al., 2013). However, the definition of connectomes by reconstruction of electron micrographic (EM) images remains an exceptionally tedious process and limits the ability to analyze the connectome of a nervous system under different conditions. This is well illustrated by the fact that to this day the connectome of only a single adult *C. elegans* hermaphrodite has been fully reconstructed. The significance of this shortcoming is illustrated by the well-recognized notion that the many dynamic aspects of brain function, from the modulation of behavior to memory formation, rely on the alterations of synaptic circuitry (LeDoux, 2014; McEwen, 2010; Takeuchi et al., 2014; Wiesel and Hubel, 1965). A nervous system-wide appreciation of the plasticity of neuronal connectomes is, however, still lacking and is the main subject of this study.

Connectomes are defined by two types of synapses, chemical and electrical synapses (gap junctions). While the role of chemical synapses in nervous system function has been widely studied, electrical synapses have received much less attention. The importance of electrical synapses is, however, well documented through genetic analysis in invertebrate and vertebrate nervous system, in which the loss of constituent components of electrical synapses result in obvious dysfunctions of the nervous system (Abrams and Scherer, 2012; Hall, 2017; Hasegawa and Turnbull, 2014; Marder et al., 2017; Song et al., 2016; White and Paul, 1999). Unfortunately, since electrical synapses are difficult to detect in currently used high-throughput EM methodologies, a map of electrical synapses (“electrical connectome”) is absent from all currently available large scale connectomes, except the *C. elegans* connectome. Within the *C. elegans* connectome, electrical synapses are wide-spread: every one of the 118 neuron classes of the *C. elegans* hermaphrodite makes electrical synapses and each class makes electrical synapses to an average number of 9.7 synaptic partners (range: 1 to 30) (Jarrell et al., 2012; White et al., 1986). Moreover, the patterns of electrical synaptic connectivity show only weak correlation with the patterns of chemical synaptic connectivity (Varshney et al., 2011).

Electrical synapses are composed of two analogous families of proteins, connexins in vertebrates (encoded by 20 distinct genes in mammals) and innexins in invertebrates (encoded by 25 genes in *C. elegans)* (Phelan and Starich, 2001; Willecke et al., 2002). Electrical synapses are formed by the assembly of either homogenous or heterogeneous combinations of connexins or innexins (Mese et al., 2007; Miller and Pereda, 2017; Phelan and Starich, 2001) (Fig. 1A). The diverse molecular composition of electrical synapses is thought to encode: (a) synaptic specificity, i.e. the choice of which neurons engage in electrical synaptic contact, based on a matching assembly of innexin proteins in connected neurons; and (b) functional diversity, i.e. electrical synapses with distinct molecular compositions are thought to have different functional properties (Baker and Macagno, 2017; Goodenough and Paul, 2009; Mese et al., 2007; Miller and Pereda, 2017; Sohl et al., 2005). The establishment of neuron-type specific patterns of innexin gene expression is therefore a likely determinant of the electrical synapse development and function.

We describe here the expression pattern of all *C. elegans* innexin genes and find that 14 innexin genes are expressed in a highly combinatorial manner in the nervous system (98 out of the 118 *C.elegans* neuron classes can be described by unique combinations of innexin genes). We discover a striking extent of plasticity of neuron-type specific expression of innexin genes in response to environmental cues (86/118 neuron classes change their combinatorial innexin code), thereby predicting a substantial rewiring of the electrical connectome. The environmental conditions that we test are adverse environmental conditions – starvation, high population density or increased temperature – that trigger the entry of *C. elegans* larvae into a diapause-arrest stage, the dauer stage (Cassada and Russell, 1975; Fielenbach and Antebi, 2008; Riddle and Albert, 1997). Honing in on a single innexin in a single neuron type, we show that the dauer-specific induction of *inx-6* in the AIB interneuron class is both necessary and sufficient to form multiple *de novo* electrical synapses. We characterize alterations of the locomotory features of dauer-stage animals and demonstrate that some of these locomotory changes, as well as some altered sensory behaviors of the dauer, are mediated through the rewiring of specific electrical synapses (including the novel INX-6-mediated electrical synapses of AIB). Lastly, we demonstrate a gene regulatory strategy by which these changes in electrical synaptic connectivity are orchestrated by the constitutive activity of cell type specific homeodomain transcription factors in conjunction with the stress-induced, cell autonomous activation of insulin/IGF-1 target DAF-16/FOXO.

## RESULTS

### A nervous system-wide molecular map of the electrical connectome

Our overall strategy to investigate the plasticity of synaptic wiring was to first establish a map of molecular determinants of synaptic connectivity and to then identify conditions under which the expression of these synaptic connectivity factors are modulated, thereby predicting changes in synaptic connectivity and circuit function. In addition, our implicit approach was to exploit the relative simplicity of the *C. elegans* nervous system to undertake such an analysis on a genome- and nervous system-wide manner. Since electrical synapses are defined by the combinatorial expression of specific innexin proteins (Fig. 1A) (Mese et al., 2007; Miller and Pereda, 2017; Phelan and Starich, 2001), we sought to precisely map the expression of all neuronal innexins.

**Fig. 1:**
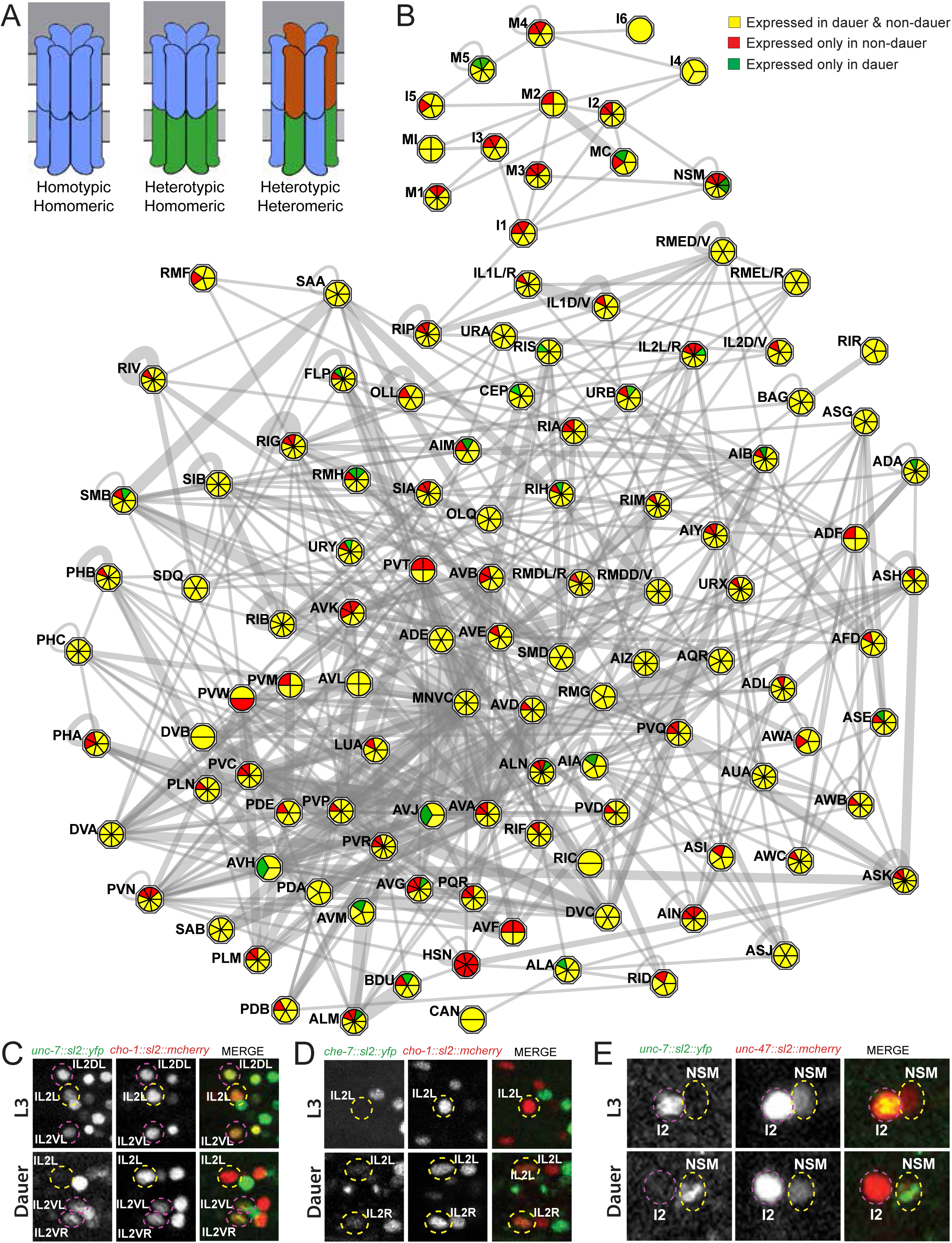
Changes in innexin gene expression in dauer. **(A)** Schematics of subunit composition in different kinds of gap junction channels. **(B)** Connectivity of the neuronal classes by electrical synapses (electrical connectome) at the adult stage, as inferred from the serial section reconstruction of electron micrographs. Neuronal classes are represented as hexagons and electrical synapses between or within different neuronal classes are shown as gray lines. Thicknesses of the lines were weighted according to the volume of each connection (measured by the total number of EM sections in which each en passant synapse was observed; as per www.wormwiring.org). The number of innexin reporter genes expressed in each neuronal class is indicated by the overlaid pie chart. Color of each section in the pie chart represents whether the reporter is expressed in both non-dauer and dauer stages (yellow) and only in non-dauer stages (red) or only in dauer stage (green). **(C-E)** Neuronal identities were determined by expression of either *otIs544[cho-1*^*fosmid*^::*sl2::mcherry::H2B]* or *otIs564[unc-47*^*fosmid*^::*sl2::mcherry::H2B]* reporters. **(C)** An sl2-based *unc-7* transcriptional fosmid reporter *(otEx7106[unc-7*^*fosmid*^::*sl2::yfp::H2B])* is expressed in dorsal IL2DL/R, ventral IL2VL/R and lateral IL2 L/R in non-dauer stages, but selectively down-regulated in lateral IL2 L/R in dauer stage. ***(D)*** *che-7* fosmid reporter *(otEx7112[che-7*^*fosmid*^::*sl2::yfp::H2B])*, which is not expressed in any IL2 neurons in non-dauer stages, is selectively turned on in lateral IL2 L/R neurons in dauer. ***(E)*** *unc-7* fosmid reporter *(otEx7106[unc-7*^*fosmid*^::*sl2::yfp::H2B])* is expressed in pharyngeal neuron I2, but not in NSM in non-dauer stages. In dauer, *unc-7* reporter expression disappeared in I2, while turned on in NSM.

A previous study using antibody staining revealed that two innexins, INX-21 and INX-22 are exclusively expressed in the gonad and thus are not included in this study (Starich et al., 2014). For *inx-2* and *inx-6*, fluorescent reporter alleles were generated using CRISPR/Cas9-mediated genome editing (see Experimental Procedures). For the remaining 21 innexin family genes, we generated fosmid-based transcriptional reporter constructs containing ∼30-40kb genomic region including the sequence of the gene of interest and ∼10-20kb flanking sequences on either side. It is thought that all the necessary *cis*-regulatory information for the endogenous gene expression is generally contained within the span of such fosmid clones. For each reporter, we introduced an SL2 trans-splicing sequence followed by an NLS and a histone (H2B)-tagged *yfp*-gene sequence right after the stop codon of the innexin gene locus (Tursun et al., 2009). These so-called “transcriptional reporters” allowed nuclear localized H2B-tagged YFP reporter protein expression independent of the membrane localized innexin protein expression, facilitating neuronal identification. Schematics of all generated transcriptional fosmid reporter clones and reporter alleles are shown in Fig. 3, **S2-3**. Three innexin genes, *inx-1, inx-10* and *inx-18,* contain alternate splice-isoforms with different 3’ end sequences. Independent SL2-based fosmid reporter constructs were generated to avoid any potential isoform specific expression bias. Among them, expression of *inx-10b* could not be faithfully identified after examining multiple independent transgenic lines and thus deemed not expressed.

**Fig. 2:**
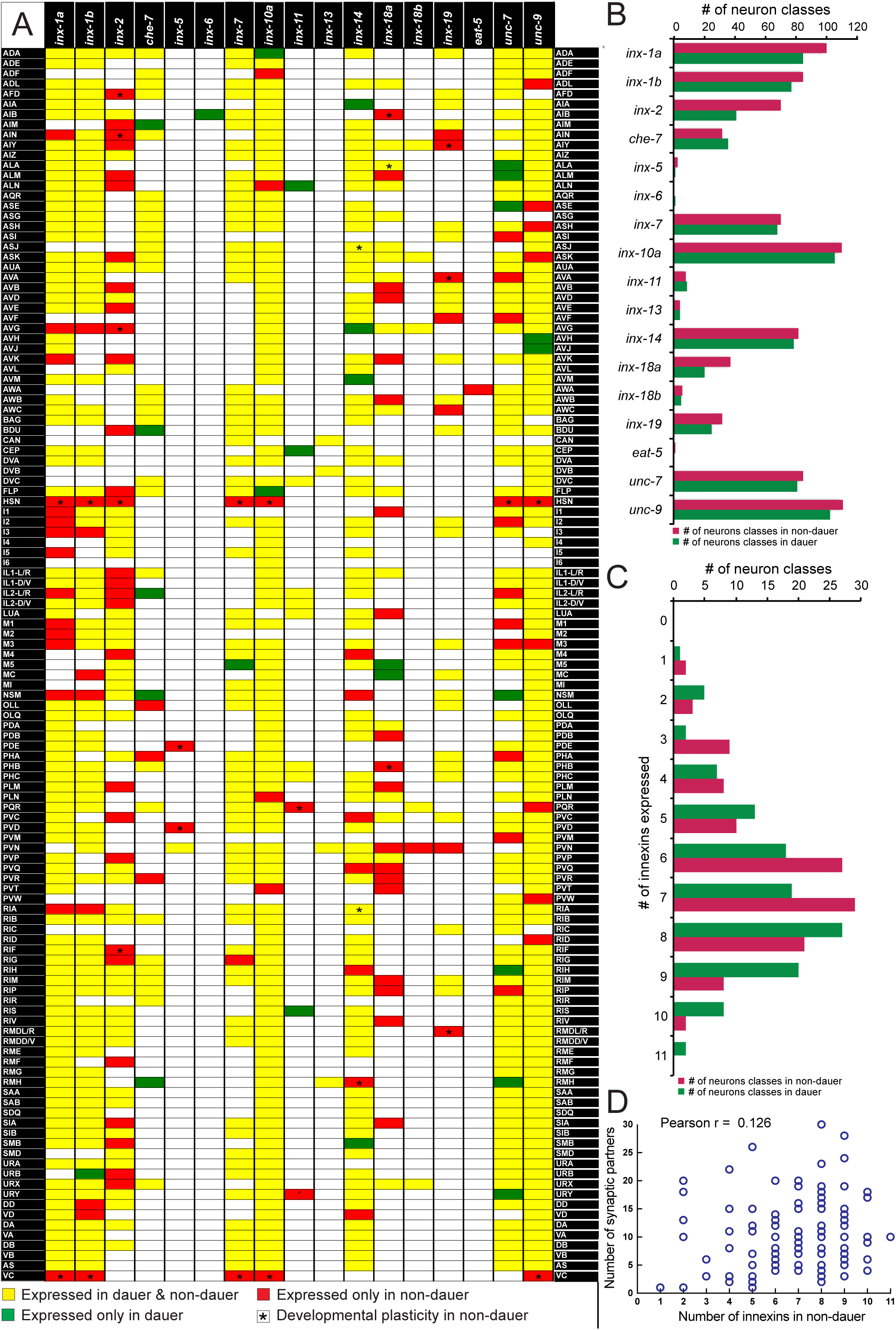
Innexin gene expression in non-dauer and dauer. **(A)** Expression of all neuronally expressed innexin genes in all neuronal classes. Color of each box in the table represents whether the innexin gene is expressed in both non-dauer and dauer stages (yellow), only in non-dauer stages (red) or only in dauer stage (green). Asterisks indicate developmental changes in innexin expression among non-dauer stages (See also Table S2 for developmental changes). We have seen differential innexin expression among 3 neurons classes (IL1, IL2 and RMD). These neuron classes were further divided in subclasses. **(B)** A graphical representation showing abundance of each innexin genes in non-dauer and dauer nervous system. **(C)** A graphical representation showing distribution of number of innexin genes expressed by neuron classes in non-dauer and dauer nervous system. **(D)** Correlation between number on innexin genes expressed by neuron classes in non-dauer stage and number of electrical synaptic targets, as inferred from the serial section reconstruction of electron micrographs (data source www.wormwiring.org) (Jarrell et al., 2012). See also Fig. S1-3, Table S1, S2.

**Fig. 3:**
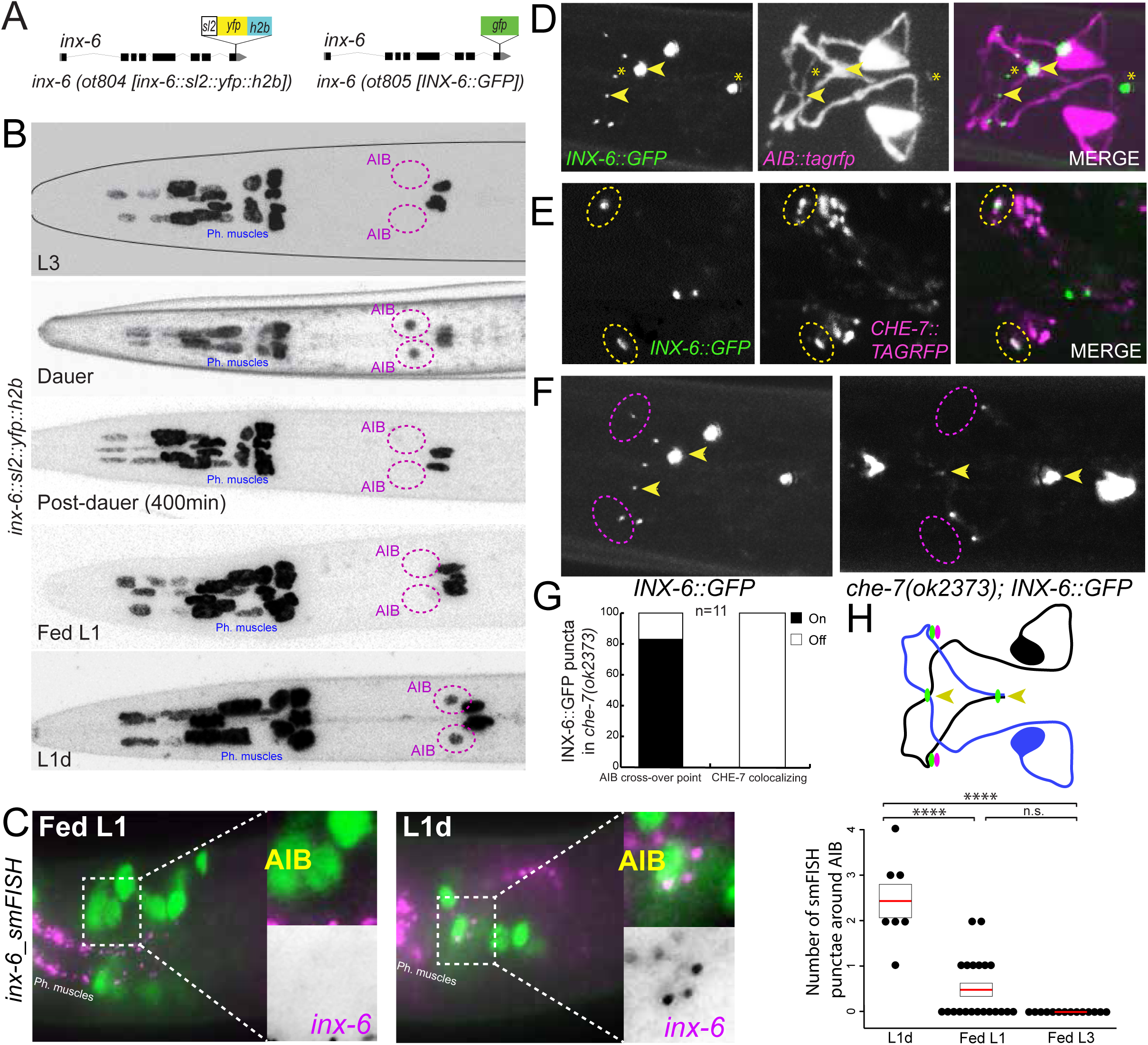
Dauer-induced expression of *inx-6* in AIB generates new gap junctions with *che-7*. **(A)** Schematics of an sl2-based *inx-6* transcriptional reporter allele, *inx-6(ot804)* and GFP-tagged *inx-6* translational reporter allele, *inx-6(ot805)* ***(B)*** *inx-6* transcriptional reporter allele, *inx-6(ot804),* is expressed in the pharyngeal muscles throughout the life of the animal. *inx-6* reporter allele is additionally expressed in AIB interneurons in dauer stage that disappears in post-dauer animals. *inx-6* allele is also expressed in AIB interneurons in L1-diapause stage (L1d). **(C)** smFISH analysis of endogenous *inx-6* mRNA expression (magenta) in Fed-L1 and L1d. AIB was marked by the *eat-4::yfp* (*otIs388[eat-4*^*fosmid*^::*sl2::yfp::H2B])* expression, (green). Inset shows enlargement of the AIB expression. Quantification also included smFISH data from Fed-L3 animals. Red line indicates the mean for each animal and black rectangles indicate S.E.M. for each condition. Wilcoxon rank-sum tests p-values (n.s. = non significant and ****p<0.0001). **(D)** Expression of a GFP tagged *inx-6* reporter allele, *inx-6(ot805),* in dauer. INX-6::GFP punctae (green) along AIB processes (magenta, visualized by *otIs643[npr-9p::tagrfp]* reporter expression) represented the location of gap junctions involving INX-6. Yellow arrowheads mark INX-6::GFP punctae at the cross-over point of AIBL and AIBR processes. Yellow asterisks mark INX-6::GFP punctae between pharyngeal muscles. **(E)** Co-localization of GFP-tagged INX-6 puncta in *inx-6(ot805)* allele (green) and TAGRFP-tagged CHE-7 punctae (magenta) (from *CHE-7*^*FOSMID*^::*TAGRFP* reporter transgene) in two of the gap junctions on AIB (yellow circle), in dauer. **(F)** INX-6::GFP punctae (pseudo colored white) that colocalized with CHE-7 (magenta circle) were lost in *che-7(ok2373); inx-6(ot805)* mutant dauers. INX-6::GFP puncta at the cross-over points of AIBL and AIBR processes, remained unaffected. Quantification of INX-6::GFP punctae on AIB in *che-7(ok2373)* dauers. **(G)** Quantification of INX-6::GFP punctae on AIB in *che-7(ok2373)* mutant dauers. **(H)** Schematic representation of INX-6 punctae (green circles) on AIB processes. Yellow arrowheads mark INX-6::GFP punctae at the cross-over points of AIBL and AIBR processes. CHE-7::TAGRFP punctae (magenta circles) localizes with two INX-6::GFP punctae away from the cross-over points.

We found that 14 innexin genes are expressed in the nervous system of larval and adult stage animals (Fig. 1, 2, **S2-3)**. Ten of the neuronally expressed genes, *inx-1, inx-2, che-7, inx-7, inx-10, inx-14, inx-18, inx-19, unc-7* and *unc-9,* were selectively (i.e. non-panneuronally) expressed in many (>30) neuron classes. *inx-5, inx-11* and *inx-13* were expressed in a more restricted manner in the nervous system (<10 neuron classes) and one innexin, *eat-5*, was uniquely expressed in a single neuron class. We also identified distinct expression patterns for specific splice isoforms of *inx-1* and *inx-18* (Fig. 2). While both *inx-1a* and *inx-1b* were expressed broadly in the nervous system, 15 neuronal classes expressed only one of the two isoforms. On the other hand, *inx-18a* and *inx-18b* showed strikingly different expression patterns. While, *inx-18a* was expressed broadly (in 37 distinct neuron classes), *inx-18b* expression was restricted to only six neuronal classes.

In total, each neuron class in the nervous system expressed at least one innexin gene. Most of the neuron classes expressed multiple innexin genes, while the maximum number of innexins expressed by a neuron class is 11 (Fig. 2). There is very weak correlation between the number of innexin genes expressed in a particular neuron class and the number of unique electrical synaptic partners it has (Pearson correlation coefficient 0.126) (Fig. 2D). We found 98 different combinations of innexin expression among neuron classes, hence, a striking 83% (98/118) of all *C. elegans* neuron classes are defined by unique combinations of innexin expression (Fig. 2).

The expression pattern of most innexins is stable throughout the larval and adult stage. However, *inx-2, inx-5, inx-11, inx-14, inx-18a* and *inx-19* showed dynamic expression during L2 to adult development in 16 distinct neuronal subclasses **(Table S2)**. HSN and VC neurons start to express their respective innexins starting at the L4 stage when other neuronal features of these neurons mature (Desai et al., 1988; Li and Chalfie, 1990; Pereira et al., 2015). Lastly, all innexin genes were expressed in one or more non-neuronal cell types (Fig. S2, S3; **Table S1**). The expression patterns that we describe have very little overlap with the innexin expression previously reported using very small (1kb or smaller) promoter fusions (Altun et al., 2009), a likely testament of the complexity of *cis*-regulatory control elements of the innexin loci, not captured in the previous study.

### Widespread changes in the neuron type-specificity of innexin gene expression in dauer stage animals

Since entry into the dauer stage of *C. elegans* entails a substantial remodeling of behavior (Cassada and Russell, 1975; Gaglia and Kenyon, 2009; Hallem et al., 2011a; Lee et al., 2012) (more below), we reasoned that the connectome of the dauer might also be substantially distinct. Since the presence or absence of innexins in a cell are expected to have a profound impact on synaptic partner choice and synaptic properties (Baker and Macagno, 2017; Goodenough and Paul, 2009; Mese et al., 2007; Miller and Pereda, 2017; Sohl et al., 2005), we investigated the expression patterns of innexins in the dauer stage as proxy for the electrical synaptic connectivity. We found that 11 of the 14 innexin genes normally expressed in the nervous system in larval and adult stages change their expression in a strikingly neuron-specific manner (Fig. 1, 2, **S1)**. This includes one innexin gene, *eat-*5, which is expressed in the nervous system only in non-dauer stage (i.e. its exclusive expression in AWA is turned off upon entry into the dauer stage). In striking contrast, another innexin gene, *inx-*6, is expressed only in the dauer nervous system (i.e., it is turned on exclusively in the AIB neuron class in dauer animals). Overall, we observed expression changes within 86 of the 118 neuron classes in the dauer stage (Fig. 1, 2, **S1-2)**. The maximum number of changes in innexins expression in a neuron class is three (Fig. 1, 2, **S1)**. In line with their immature state at the dauer stage, HSN and VC neurons showed no innexin expression in dauer. This plasticity in expression was observed in sensory, motor and interneuron classes.

Several aspects of the innexin expression plasticity are of particular note. Firstly, expression of distinct innexin genes could be altered in an opposing manner in a particular neuron class when animals go into dauer arrest (Fig. 1, 2). For example, in dauer stage *unc-7* expression was downregulated in lateral IL2 neurons (IL2L/R subclass), while *che-7* expression was simultaneously upregulated in the same subclass (Fig. 1C,D). Secondly, expression of a particular innexin gene could change in opposing manner in distinct neuronal subclasses during dauer molt (Fig. 1, 2). For example, in dauer, *unc-7* expression was downregulated in pharyngeal motorneuron I2, while simultaneously upregulated in another pharyngeal motorneuron NSM (Fig. 1E). Thirdly, expression of specific splice isoforms of a particular innexin gene could be independently altered in distinct neuronal subclasses during dauer molt. For example, *inx-1a* expression was specifically downregulated in IL2L/R subclass in dauer, while *inx-1b* expression remained unaltered (Fig. 1, 2). Similarly, expression of only the *inx-18a* isoform was turned on in pharyngeal M5 and MC neurons in dauer stage animals (Fig. 1, 2).

We conclude that the nervous system of *C. elegans* undergoes a widespread remodeling in the expression pattern of innexin genes. These changes in expression can be expected to have two consequences – changes in synaptic partners, prompted by the loss or gain of specific innexin genes, or changes in the signaling properties of the existing electrical synapses through changes in synaptic composition.

### Formation of INX-6-containing electrical synapses in AIB interneurons in dauer stage animals

*inx-6* reporter fusions were previously shown to be exclusively expressed in the pharynx (corpus pharyngeal muscles and in the marginal cells) throughout the life of the animal (Li et al., 2003). We confirmed this expression using both the transcriptional *inx-6* reporter allele *ot804* (generated by CRISPR/Cas9-mediated insertion of the *SL2::NLS::yfp::H2B* cassette) and the fosmid-based reporter transgene *otIs473,* respectively (Fig. 3A). As the animal molt into the dauer state, the *inx-6* transcriptional CRISPR reporter allele, *inx-6(ot804),* was additionally turned on in a single pair of interneurons, AIB (Fig. 3B). This dauer-specific expression of *inx-6* was reversible and disappeared from AIB when dauer animals resume development under favorable conditions (Fig. 3B). *inx-6* expression was also turned on in AIB during L1-diapause (L1d), which was reversible and disappeared upon feeding (Fed-L1) (Fig. 3B). To independently assess the *inx-6* expression dynamics in AIB, single molecule fluorescence in situ hybridization (smFISH) was performed to determine the endogenous *inx-6* mRNA expression. smFISH analysis showed the *inx-6* mRNA expression in pharyngeal muscles in all stages analyzed (Fig. 3C). Consistent with our reporter allele expression, smFISH analysis confirmed the expression of *inx-6* mRNA in AIB during L1d that was downregulated in Fed-L1 (Fig. 3C). Similarly, *inx-6* mRNA could not be detected in AIB at L3 stage (Fig. 3C).

To determine whether the *inx-6* expression in AIB during dauer resulted in punctate electrical synaptic contacts, we additionally generated a GFP protein-fusion allele of *inx-6* by tagging the endogenous locus with *gfp,* using CRISPR/Cas9 (*ot805* allele) (Fig. 3A). This allele did not show the lethality associated with the loss of *inx-6* function (Li et al., 2003), suggesting that *gfp-*tagging did not affect the *inx-6* function. In *inx-6(ot805)* animals, punctate GFP expression was observed in the pharyngeal muscles in all stages, while in dauer animals distinct GFP punctae were localized along the AIB processes in positions that were stereotypic across different animals (Fig. 3D,H). Among the AIB-associated INX-6::GFP punctae, two were localized at the crossover points of AIBL and AIBR processes (Fig. 3D,H). INX-6 punctae were also observed in similar positions on AIB processes during L1d (discussed later). These results suggest that the expression of an innexin family gene, *inx-6,* is dynamically regulated in the dauer nervous system and leads to the formation of either novel synaptic contacts or changes the composition of already existing AIB electrical synapses (made with six different synaptic partners according to the electron micrographic analysis) (White et al., 1986).

### INX-6 expressed in AIB functions with CHE-7 in BAG to form new electrical synapses in dauer stage animals

We next set out to analyze the INX-6 punctae formed in the AIB neurons in dauer. We first examined the strong INX-6 punctae observed at the cross-over point of the two bilaterally symmetric AIB processes, suggesting the formation of electrical synapses between these two neurons (such auto-synapses are not observed in non-dauer animals) (Jarrell et al., 2012; White et al., 1986). To test this hypothesis, we surgically removed one of the bilateral AIB neurons and observed that such ablation leads to selective loss of the INX-6 punctae at the point where the AIB neuronal processes normally cross (Fig. 4A). This result suggests that the bilateral AIB neurons may electrically couple to one another selectively in dauer-stage animals.

**Fig. 4:**
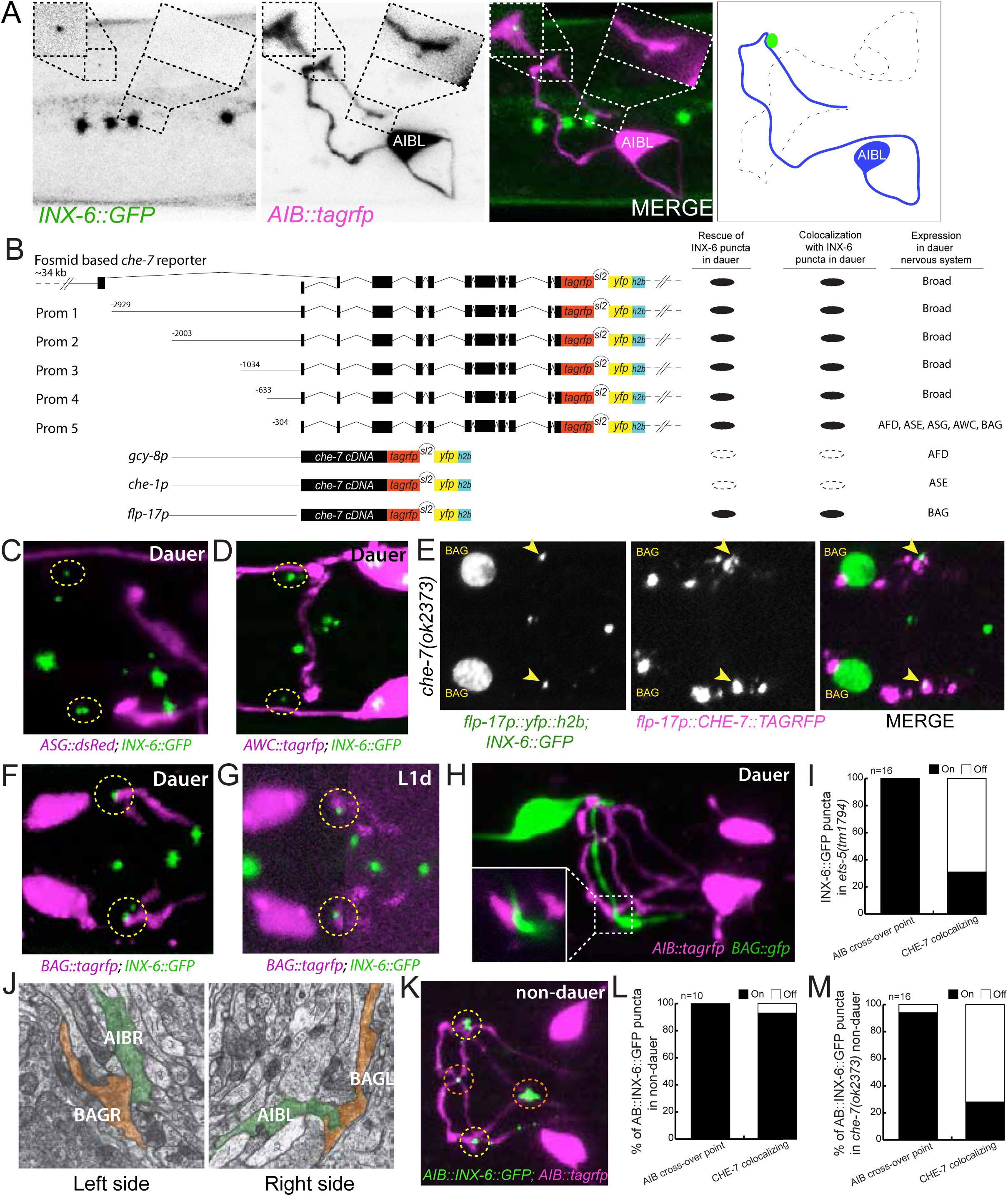
*inx-6* and *che-7* forms gap junction between AIB and BAG. **(A)** Expression of INX-6::GFP punctae (green) on AIBL (magenta) in *daf-7(e1372)* dauer animals (*inx-6(ot805); daf-7(e1372); otIs643[npr-9p::tagrfp])*, where AIBR was ablated using laser micro-beam. In absence of AIBR, INX-6::GFP punctae that co-localized with CHE-7 (left box) remains intact, while INX-6::GFP punctae at the crossover point of AIBL and AIBR processes (right box) disappear. Right panel shows the schematic representation of the effect of AIBR-ablation on INX-6 punctae on the remaining AIBL neuron. Green circle represents the CHE-7-associated INX-6 punctae at the place of AIB-BAG electrical synapses. **(B)** Schematics of 5’ cis-regulatory element analysis of *che-7* and reporter transgenes used for cell-specific *che-7* expression. Transgenic lines were created in *che-7(ok2373); inx-6(ot805)* background. 2 to 3 independent transgenic reporter lines were scored (n≥10 per line). Lines commonly showed very similar expressivity and penetrance. Average results for all transgenic lines are shown. Strains are listed in Table S5. **(C-D)** INX-6::GFP punctae (green) that co-localize with CHE-7 in dauer (yellow dotted circles) do not co-localize with ASG (*inx-6(ot805);oyIs47[ops-1p::dsRed2])* and AWC (*inx-6 (ot805);otIs263[ceh-32p2::tagrfp]* axons (magenta). **(E)** Using the *flp-17p::che-7::tagrfp::sl2::yfp::H2B* transgene labels the nuclei of BAG neurons (green) and expresses CHE-7::TAGRFP fusion protein (magenta) only in BAG neurons. In *che-7(ok2373)* dauer, expression of CHE-7::TAGRFP only in BAG neurons rescues CHE-7 associated INX-6::GFP punctae (green) that also shows colocalization with CHE-7::TAGRFP (yellow arrowheads). (*inx-6(ot805); che-7(ok2373); otEx2487(flp-17p::che-7::tagrfp::sl2:: yfp::h2b])* **(F)** INX-6::GFP punctae (green) in dauer co-localize with BAG axon (magenta). (*inx-6(ot805); otEx7230[ets-5p::tagrfp])* **(G)** INX-6::GFP punctae (green) in L1d co-localize with BAG axon (magenta). For BAG-axon image clarity, a projection of non-continuous z-sections was shown. (*inx-6(ot805); otEx7230[ets-5p::tagrfp])* **(H)** BAG axon (green) comes in contact with AIB process (magenta) at similar positions where INX-6::GFP punctae co-localize with BAG axon. Inset shows enlargement of the contact point. (*otIs643[npr-9p::tagrfp]; otEx7230[ets-5p::tagrfp])* **(I)** Quantification of INX-6::GFP punctae on AIB in *ets-5(tm1794)* mutant dauers. **(J)** TEM prints from wild-type adult hermaphrodite ‘N2U’ showing adjacency of BAG (pseudo colored in red) and AIB (pseudo colored in green) neuronal processes at the site where dauer specific INX-6-CHE-7 electrical synapses are formed. These images were collected in MRC/LMB and annotated images were obtained from www.wormimage.org, courtesy of David Hall. Prints shown here are, Left: N2U_094 and right: N2U_116. **(K)** Ectopic expression of INX-6::GFP specifically in AIB (*otTi19)* in non-dauer stage (L3) shows INX-6::GFP punctae (green) along the AIB processes (magenta). Red circles mark INX-6::GFP punctae at the crossover point of AIBL and AIBR processes. Yellow circles mark INX-6::GFP punctae at the site where dauer specific INX-6-CHE-7 electrical synapses are formed. (*otTi19 (Si[Pnpr-9::INX-6::GFP]); otIs643[npr-9p::tagrfp])* **(L)** Quantification of INX-6::GFP punctae in non-dauer animals ectopically expressing INX-6::GFP specifically in AIB (*otTi19*). **(M)** Quantification of INX-6::GFP punctae in non-dauer *che-7(ok2373); otTi19(Si[Pnpr-9::INX-6::GFP])* animals.

To characterize the other highly stereotyped INX-6 punctae along the AIB axons, we set out to identify innexins that may partner with INX-6 in *trans* across an electrical synapse. We based our search on two criteria: (1) the punctae formed by the partner innexin must colocalize with the INX-6 punctae on AIB and (2) formation of the INX-6 punctae must depend on the presence of such partner innexin, as is the case for other heterotypic gap junctions (Starich et al., 2009). In line with these criteria, we found that (1) TagRFP-tagged CHE-7 protein expressed from a fosmid-based reporter construct, co-localized with the INX-6 punctae in dauer stage animals at two distinct positions along the AIBL and AIBR processes, away from the crossover points (Fig. 3E,H) and that (2) loss of *che-7* results in specific loss of INX-6 punctae only at these distinct positions, but not at the AIBL-AIBR crossover points (Fig. 3F,G). Altogether, these results suggest that CHE-7 acts as the INX-6 partner to form electrical synapses specifically in AIB neurons during the dauer stage. Moreover, *che-7,* although broadly expressed, is not expressed in AIB in both non-dauer and dauer stages (Fig. 2). This suggests that CHE-7 must be expressed in the putative AIB synaptic partner neurons and interacts with INX-6 in *trans* across the synapse.

To identify the putative AIB synaptic partner neurons, we reasoned that *che-7* expression in such a partner neuron should be (a) sufficient to rescue the loss of INX-6 electric synapse punctae in the *che-7* mutant dauer animals and (b) should co-localize with the INX-6 punctae. To address this question, we generated a fosmid-based bi-cistronic reporter construct where *che-7* locus was tagged at the 3’ end with *tagrfp* followed by an SL2 trans-splicing sequence separated H2B-tagged *yfp* cassette (Fig. 4B). This reporter allowed independent visualization of the membrane-localized TagRFP-tagged CHE-7, while simultaneously allowed identification of the *che-7* expressing neurons based on the nuclear-localized YFP (Fig. 3B). Corroborating our premise, expression of this fosmid-based bi-cistronic *che-7* reporter in *che-7* mutant dauers was sufficient to rescue the two CHE-7-associated INX-6 punctae and showed colocalization of the TagRFP-tagged CHE-7 with GFP-tagged INX-6. Mimicking the transcriptional *che-7* fosmid reporter, this bi-cistronic fosmid reporter also showed broad expression in the nervous system (Fig. 2A, 4B, **S2)**. To narrow down to the putative CHE-7-expressing AIB-synaptic partner, we further divided the *cis*-regulatory region driving the expression of the bi-cistronic *che-7* reporter cassette into gradually smaller elements. A 304bp minimal *cis*-regulatory region that was expressed in five sensory neuron pairs (ASG, AWC, AFD, ASE and BAG) was sufficient to rescue the INX-6 electrical synapse punctae in *che-7* mutant dauers and showed colocalization of CHE-7 with INX-6 (Fig. 4B). Through visualization of the AWC and AFD axons, we found that these processes are not in proximity to the INX-6::GFP punctae (Fig. 4C,D). Single neuron specific expression of the bi-cistronic *che-7* reporter cassette showed that *che-7* expression in the BAG neuron alone was sufficient to rescue the INX-6 punctae in *che-7* mutant dauer and showed colocalization of CHE-7::TagRFP with INX-6::GFP (Fig. 4E). When crossed with the BAG neuron specific axon tracer, INX-6::GFP punctae colocalized with BAG axons in dauer stage animals (Fig. 4F) as well as in L1d animals where INX-6::GFP is also induced in AIB (Fig. 4G). BAG and AIB projections also came into contact with each other at similar positions (Fig. 4H). Moreover, the two CHE-7-associated INX-6 punctae were lost in dauers mutant for *ets-5,* an ETS domain transcription factor required for the BAG neuron differentiation (Guillermin et al., 2011), while the INX-6::GFP punctae at the AIBL-AIBR crossover points remained unaffected (Fig. 4I).

At the adult stage AIB and BAG do not form any electrical synapses, as inferred from the serial section reconstruction of electron micrographs (White et al., 1986). Analyzing existing serial section electron micrographs of adult animals, we found that the AIB and BAG neuronal processes are directly adjacent to each other at positions similar to the site of INX-6-mediated electrical synapse formation in dauer (Fig. 4J), indicating that the absence of AIB and BAG electrical synapses in non-dauer animals may solely be the result of the absence of INX-6 expression. We indeed found that ectopic expression of *inx-6* in AIB of non-dauer animals is sufficient to form CHE-7-dependent puncta at the BAG-AIB contact points (Fig. 4K-M). AIB-specific ectopic expression of INX-6 in non-dauer animals also led to electrical synapse formation at the AIBL-AIBR crossover points (Fig. 4K,L). These results together suggest that the regulation of *inx-*6 expression in AIB is both necessary and sufficient for the induction of BAG-AIB electrical synapses.

### Dauer animals display distinctive locomotory behaviors

Having defined the system wide changes in innexin expression as well as specific changes of the electrical connectome in the dauer stage, we sought to define the physiological relevance of these changes. Several changes in the behavior of dauer animals have been described before. For example, dauers exhibit prolonged bouts of spontaneous quiescence (Gaglia and Kenyon, 2009), a strong reduction in pharyngeal pumping rate (Cassada and Russell, 1975) and nictation behavior, where an animal stands vertically on its tail and waves its head (Cassada and Russell, 1975; Lee et al., 2012). We sought to further expand the known repertoire of altered behaviors of dauers by quantitatively assessing the locomotory patterns of dauer and non-dauer animals using an automated Wormtracker 2.0 system (Yemini et al., 2013). Specifically, we compared three different stages and states of the worm: (1) Wild-type laboratory Bristol N2 wild-type strain animals in the dauer stage. The decision to enter this state is made during the L1 stage (Schaedel et al., 2012) and developmental stage wise is equivalent to L3 stage animals that have not experienced adverse conditions. We induced dauer arrest under standardized starvation, crowding and high-temperature conditions. (2) Animals in L3 larval stage (Fed-L3). (3) To assess the impact of starvation alone, we also analyzed L3 stage animals after a 4h-8h interval with no food (starved-L3). As dauer animals are thinner, more elongated and contain a different cuticular structure compared to fed- or starved-L3 animals (Cassada and Russell, 1975), some of the behavioral differences measured by the Wormtracker (e.g. wavelength) that are likely to be influenced by such physical differences were ignored and we instead focused on behavioral differences that can more readily be attributed to information processing in the nervous system. To minimize any potential bias arising due to dauer stage-specific prolonged bouts of spontaneous pausing (Gaglia and Kenyon, 2009), we only selected animals that were actively moving at the beginning of the assay.

Principal component analysis (PCA) of 195 locomotory behavioral features obtained from the Wormtracker revealed that the behaviors of dauer, fed- and starved-L3 animals could be clustered in distinct, separate groups (Fig. 5A,B; **Table S3**). Upon further examination of the individual behavioral features, we found three different patterns of changes (Fig. 5; **Table S4)**. In pattern #1, dauers differ from both fed-L3s and starved-L3s. Examples include head bend amplitude (Fig. 5C,D) and omega turn behavior, where animals change direction by attaining an intermediate omega symbol (O) like posture, and is often associated with the initiation of backing (Fig. 5C,E). Dauers performed omega turns less frequently than both fed- and starved-L3s (Fig. 5E). In pattern #2, both starved-L3s and dauers differ from fed-L3s in a similar manner. Examples include coiling and backward motion behaviors (Fig. 5C,F,G). We found that both starved-L3s and dauers performed coiling much less frequently and showed much less backward motion (measured as backward motion time ratio = total time spent moving backward/total time of the assay). Finally, in pattern #3, we observed both dauers and fed-L3s differ from starved-L3s in a similar manner. Examples include foraging, pausing and forward motion behaviors (Fig. 5C,H-K). The behavioral differences discussed in pattern #3 indicate that the prolonged starvation-induced habituation may influence dauers to behave as fed-L3s for certain aspects of the dauer behavior, such as food search behavior.

**Fig. 5:**
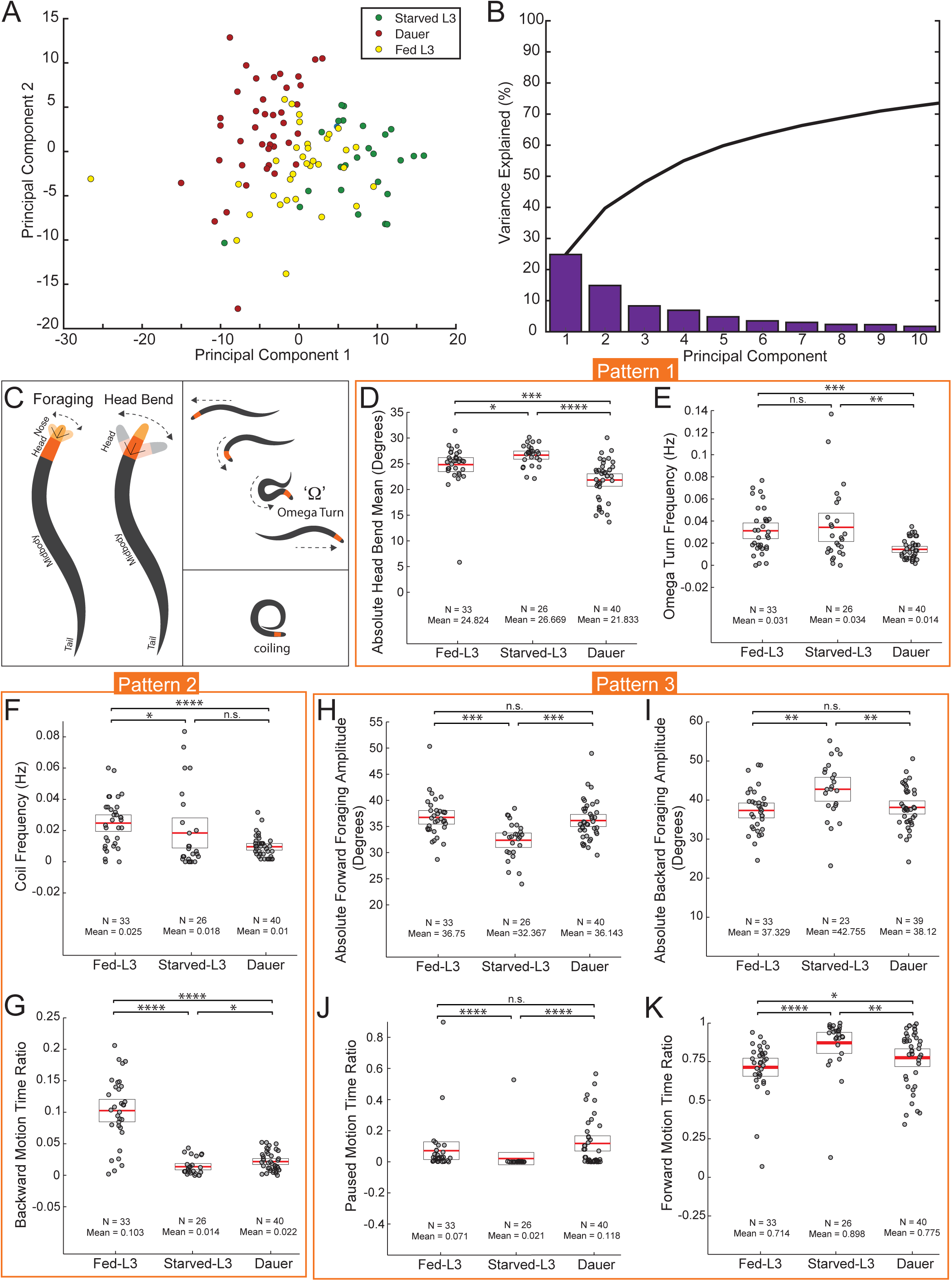
Locomotory behavior is remodeled in dauer. **(A)** Principal Component Analysis of dauer (red), fed-L3 (yellow) and starved-L3 (green) animals based on 195 locomotory behavior feature data (Listed in table S3). Circles represent individual animals (n^dauer^ = 40, n^Fed-L3^ = 33, n^starved-L3^ = 26). Component 1 and 2 account for ∼40% of the variation in the locomotion behaviors. **(B)** Variance explained by first ten PCs. Black line represents cumulative variance explained. **(C)** Schematics of the foraging behavior (nose bend), head bend, reversal through an omega turn and coiling. **(D-K)** Worm locomotion assay using a single worm tracker, Wormtracker 2.0 system (see Methods for detailed description of behavioral assays). Each circle represents the experimental mean of a single animal for the corresponding behavioral assay. Red lines indicate the mean of means and black rectangles indicate S.E.M. for each assay. q-values for each comparison (n.s. = non significant, *q<0.05, **q<0.01, ***q<0.001, ****q<0.0001) were determined by Wilcoxon rank-sum tests and False-Discovery Rate. We observed three distinct patterns of behavioral changes between the three groups of animals analyzed. In Pattern 1 (D-E), dauer behavior is different from both fed- and starved-L3 behavior. In Pattern 2 (F-G), dauer and starved-L3 behavior is different from fed-L3 behavior in a similar manner. In Pattern 3 (H-K), dauer and fed-L3 behavior is different from starved-L3 behavior in a similar manner. **(D)** Dauer animals showed significant decrease in head bend mean/amplitude compared to both fed and starved-L3 animals. Starved-L3s showed small but significant increase in head bend mean compared to fed-L3. **(E)** Dauer animals showed significant decrease in omega turn frequency compared to both fed and starved-L3 control animals, while here was no difference between fed- and starved-L3. **(F)** Dauers and starved-L3s showed significant decrease in coil frequency compared to fed-L3s. **(K)** Dauers and starved-L3s showed significant decrease in backward motion time ratio compared to fed-L3s. **(H and I)** Dauer and fed-L3 animals showed similar significant increase or decrease in foraging amplitude while moving forward or backward, respectively, compared to starved-L3 animals. **(J)** Dauer and fed-L3 animals showed similar significant increase in pause motion time ratio compared to starved-L3 animals. **(K)** Dauer and fed-L3 animals showed similar significant decrease in forward motion time ratio compared to starved-L3s. See also Table S3 and S4.

### *inx-6* and *che-7* affect locomotory and chemotaxis behavior specifically in dauer

To assess whether formation of the dauer-specific *de novo* INX-6 electrical synapses attributes new functions to AIB, we first assessed the consequences of AIB removal through caspase-driven genetic ablation of AIB (Wang et al., 2017). In fed and starved non-dauer animals, consistent with the previous reports (Gray et al., 2005), AIB-ablation severely affected backward locomotion and subsequently enhanced forward locomotion (Fig. S4E-J). However, AIB acquires novel functions in dauer locomotory behavior. As shown above in Fig. 5, dauers showed a distinctive extent of pausing, forward and backward locomotion compared to non-dauer L3-staged animals and ablation of AIB affected all three of these behaviors (Fig. 6A-C **and S4A,C)**.

**Fig. 6:**
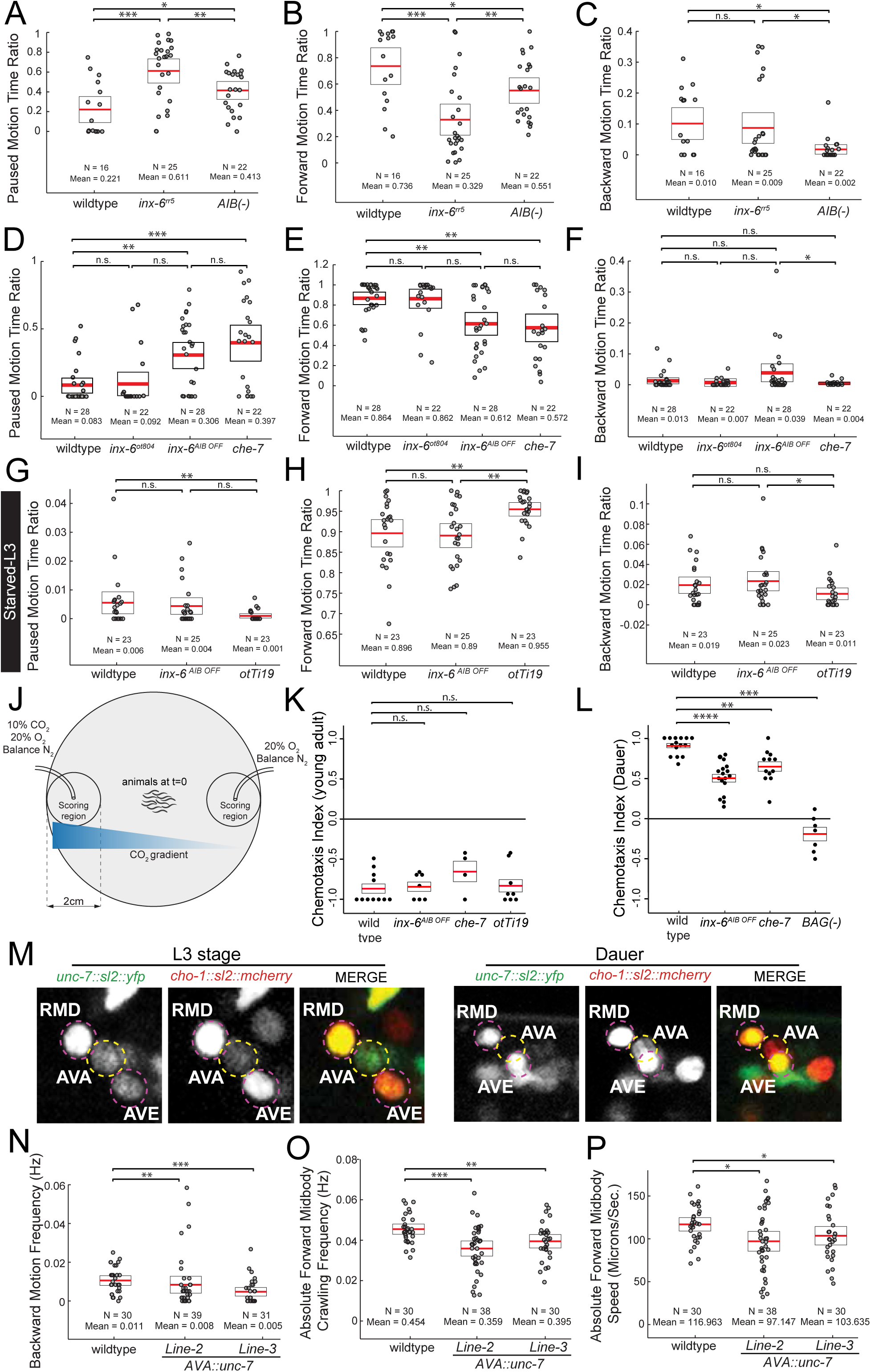
Loss and gain of innexin expression in dauers affect locomotory behavior and CO_2_-attraction behavior. **(A-I)** Worm locomotion assay using Wormtracker 2.0 (see Methods for detailed description). Each circle represents the experimental mean of a single animal. Red lines indicate the mean of means and black rectangles indicate S.E.M. for each assay. Wilcoxon rank-sum tests p-values for each comparison: n.s. = non significant, *p<0.05, **p<0.01, ***p<0.001, ****p<0.0001. Behavioral feature time ratio = (total time spent executing particular behavior/total assay time). **(A-F)** Locomotion of dauer animals. **(A-C)** Dauers with a temperature sensitive allele of *inx-6, inx-6(rr5),* exposed to restrictive temperature for >24h and AIB-ablated dauers (transgenic animals *peIs578[npr-9p::casp1, npr-9p::venus, unc-122p::mCherry]*, expressing caspase specifically in AIB neurons (Wang et al., 2017) show significant increase in pause motion time ratio and decrease in forward motion time ratio compared to N2 wild-type dauers. Only AIB-ablated dauers, but not *inx-6* mutant dauers show significant reduction in backward motion time ratio. **(D-F)** Dauers with *inx-6* transcriptional reporter allele, *inx-6*^*ot804*^, do not affect pause-, forward- and backward motion time ratio. Dauers that specifically lack *inx-6* expression in AIB, *inx-6*^*AIB*^*OFF* allele (see Fig. 8) and *che-7(ok2373)* allele show significant increase in pause motion time ratio and decrease in forward motion time ratio, but show no effect on backward motion time ratio. **(G-I)** In starved-L3 stage, *inx-6*^*AIB*^ ^*OFF*^ animals show no significant difference in pause-, forward- and backward motion time ratio. Ectopic expression of INX-6::GFP specifically in AIB in starved-L3 animals (*otTi19 (Si[Pnpr-9::INX-6::GFP])* shows decrease in pause motion time ratio and increase in forward motion time ratio compared to N2 wild-type animals. Ectopic expression of INX-6::GFP shows no effect on backward motion time ratio in starved-L3 animals. **(J)** Schematic of CO_2_-chemotaxis assay (See Methods for detailed description of CO_2_-chemotaxis assay). **(K-L)** Each circle represents chemotaxis index calculated from a single assay. Red lines indicate the mean and rectangles indicate S.E.M. Wilcoxon rank-sum tests p-values for each comparison: n.s. = non significant, *p<0.05, **p<0.01, ***p<0.001, ****p<0.0001. ***(K)*** *inx-6*^*AIB*^ ^*OFF*^ or *che-7(ok2373)* mutant young adults do not affect CO_2_-repulsion in young adults. Ectopic expression of INX-6::GFP in AIB (*otTi19 (Si[Pnpr-9::INX-6::GFP])* in young adults do not affect CO_2_-repulsion. ***(L)*** *inx-6*^*AIB*^ ^*OFF*^ and *che-7(ok2373)* dauers show significant reduction in CO_2_-attraction. Dauers with ablated BAG neurons (By Caspase expression specifically in BAG neurons, *kyIs536; kyIs538*) do not chemotax to CO_2_ gradient. See also Fig. S4. **(M)** Expression of the fosmid-based *unc-7* transcriptional reporter *(otEx7106[unc-7*^*fosmid*^::*sl2:: yfp::H2B])* is lost in AVA interneurons as the animals molt into dauer state, while continue to be expressed in RMD and AVE. *otIs544[cho-1*^*fosmid*^::*sl2::mcherry::H2B]* reporter expression was used for neuronal identification. **(N-P)** Locomotion of two independent transgenic lines where *unc-7* is ectopically expressed in AVA in dauers (Transgenic Line-2: *otEx7250 [flp-18p::unc-7::sl2::yfp::H2B]* and Line-3: *otEx7251 [flp-18p::unc-7::sl2::yfp::H2B]*). See also Fig. S4.

To address the role of *inx-6* in the functional plasticity of AIB, we tested the locomotion of *inx-6* mutant dauers. Since *inx-*6 loss-of function mutation is L1 lethal, we used a temperature-sensitive allele of *inx-6, inx-6(rr5),* for the dauer locomotion assay (Li et al., 2003). In addition, to assess the role of INX-6 specifically in AIB, we used CRISPR/Cas9-mediated genome editing to generate an *inx-6*^*AIB*^ ^*OFF*^ allele that removes *inx-6* expression exclusively in the AIB neurons, leaving the pharyngeal muscle expressions intact (discussed later in detail). Dauers mutant with either of the *inx-6* alleles, showed pausing and forward locomotion defects similar to the AIB-ablated dauers, however, did not affect the backward locomotion, which was also affected by AIB-ablation (Fig. 6A-F). Dauers mutant for *che-*7, the innexin partner of *inx-*6, expressed in BAG, showed exactly the same locomotory defects (Fig. 6A-F). Consistent with the observation that *inx-6* is only expressed in the nervous system at the dauer stage, *inx-6* mutations did not affect the pausing or forward locomotion in fed or starved non-dauer animals (Fig. 6G-I, **S4E-J)**.

Since ectopic expression of *inx-6* in AIB was sufficient to form electrical synapses in non-dauer animals (Fig. 4K), we next asked whether these ectopic electrical synapses were also sufficient to alter non-dauer locomotion. We indeed found that the AIB-specific ectopic *inx-6* expression reduced pausing and enhanced forward locomotion in non-dauer animals Fig. 6G-I), indicating that the induction of an innexin gene expression in a specific neuronal subclass is both necessary and sufficient to modify the electrical connectome and is functionally relevant.

We tested additional functions of *inx-6,* now in the context of processing BAG-dependent sensory responses. *C. elegans* in non-dauer stage displays a robust chemotactic avoidance response when subjected to a CO_2_-gradient, which is sensed mainly through the BAG neuron (Hallem et al., 2011b). In contrast, dauers are attracted to high concentrations of CO_2_ in a BAG-dependent manner (Hallem et al., 2011a). Based on the findings described above, we asked whether the dauer-specific, CHE-7/INX-6 dependent BAG-AIB synapse also contribute to CO_2_ chemotaxis of dauer. Indeed, loss of either *inx-6* or *che-7* reduced CO_2_ attraction in dauer, but had no effect on CO_2_ avoidance in non-dauer animals (Fig. 6J-L). We also found that the effect of *inx-6* and *che-7* mutation was specific to CO_2_ chemotaxis and did not affect the overall chemotactic ability of dauers (Fig. S4K-O). However, unlike locomotory behaviors, AIB-specific ectopic expression of *inx-6* in non-dauer stages had no effect on CO_2_ chemotaxis (Fig. 6K), suggesting a permissive, but not an instructive role of the INX-6-CHE-7 electrical synapses in the CO_2_ chemotaxis circuit.

### Downregulation of *unc-7* expression in AVA interneurons is required for normal dauer locomotion

Moving beyond the paradigm of the induction of expression of an innexin (and the ensuing establishment of new electrical synapses), we assessed the physiological consequences of the dauer-specific suppression of an innexin gene. We considered the innexin gene, *unc-*7, which is expressed, among many other neurons, in the AVA command interneurons of the *C. elegans* motor circuit, where it is required to control the balance of forward and backward locomotion (Fig. 2, **6M**) (Kawano et al., 2011; Liu et al., 2017; Starich et al., 2009). We found that *unc-7* expression was specifically downregulated in AVA in dauers (Fig. 2, **6M)**, suggesting a remodeling of the many electrical synapses that AVA normally generates (White et al., 1986). To test the functional significance of this downregulation, we ectopically maintained *unc-7* expression in AVA in dauer and asked which of the dauer-specific locomotory behavioral alterations are potentially reverted back to a more non-dauer state. Multiple distinct parameters that measure the agility of worms are increased in dauers (Fig. 5, **Table S4)** and we found that transgenic, dauer-stage animals force-expressing *unc-7* in AVA showed reduced agility, resembling more the non-dauer stage (Fig. 6N-P). Taken together, remodeling of the electrical synapses of the AVA neurons predicted upon *unc-7* downregulation is partly responsible for the remodeling of locomotory behavior of dauer animals.

### DAF-16/FoxO intersects with terminal selector function to ensure dauer-specificity of innexin expression

To understand the regulatory logic of the condition-specific innexin expression plasticity, we returned to the *inx-6* locus, analyzing its *cis*-regulatory control elements. We generated transgenic animals carrying fluorescent reporter transgenes containing various fragments of the *inx-6 cis*-regulatory region and compared their expression with the *inx-6* reporter allele, *inx-6(ot804*) and the *inx-6* fosmid reporter, *otIs473* (Fig. 7A). We identified a minimal 480bp sequence that was sufficient to recapitulate dauer-specific expression in AIB (Prom 7) (Fig. 7A). Further deletions defined a 144bp sequence containing required *cis*-regulatory information for AIB expression (Fig. 7A). Within this 144bp region, we observed a predicted binding site for a homeodomain transcription factor that is completely conserved in multiple nematode species (TAATTA). Through a systematic analysis of homeobox gene expression, we identified the paired-type homeodomain transcription factor *unc-42,* related to mammalian Prop1 (Baran et al., 1999), as being expressed in AIB throughout the life of the animal (Fig. 7C). The expression of *inx-6* reporter transgenes was lost in AIB in *unc-42* mutant background (Fig. 7B). We deleted the predicted UNC-42 binding site in the genome in the context of the *inx-6* transcriptional reporter allele, *inx-6(ot804)* (Fig. 7A). These alleles were specifically defective for *inx-*6 expression in AIB in dauer and L1d, while expression in the pharyngeal muscles remained unaffected (Fig. 7G), corroborating the requirement of UNC-42 in the spatial regulation of *inx-6*. We termed these alleles as the *inx-6*^*AIB*^ ^*OFF*^ alleles that we had already mentioned above.

**Fig. 7:**
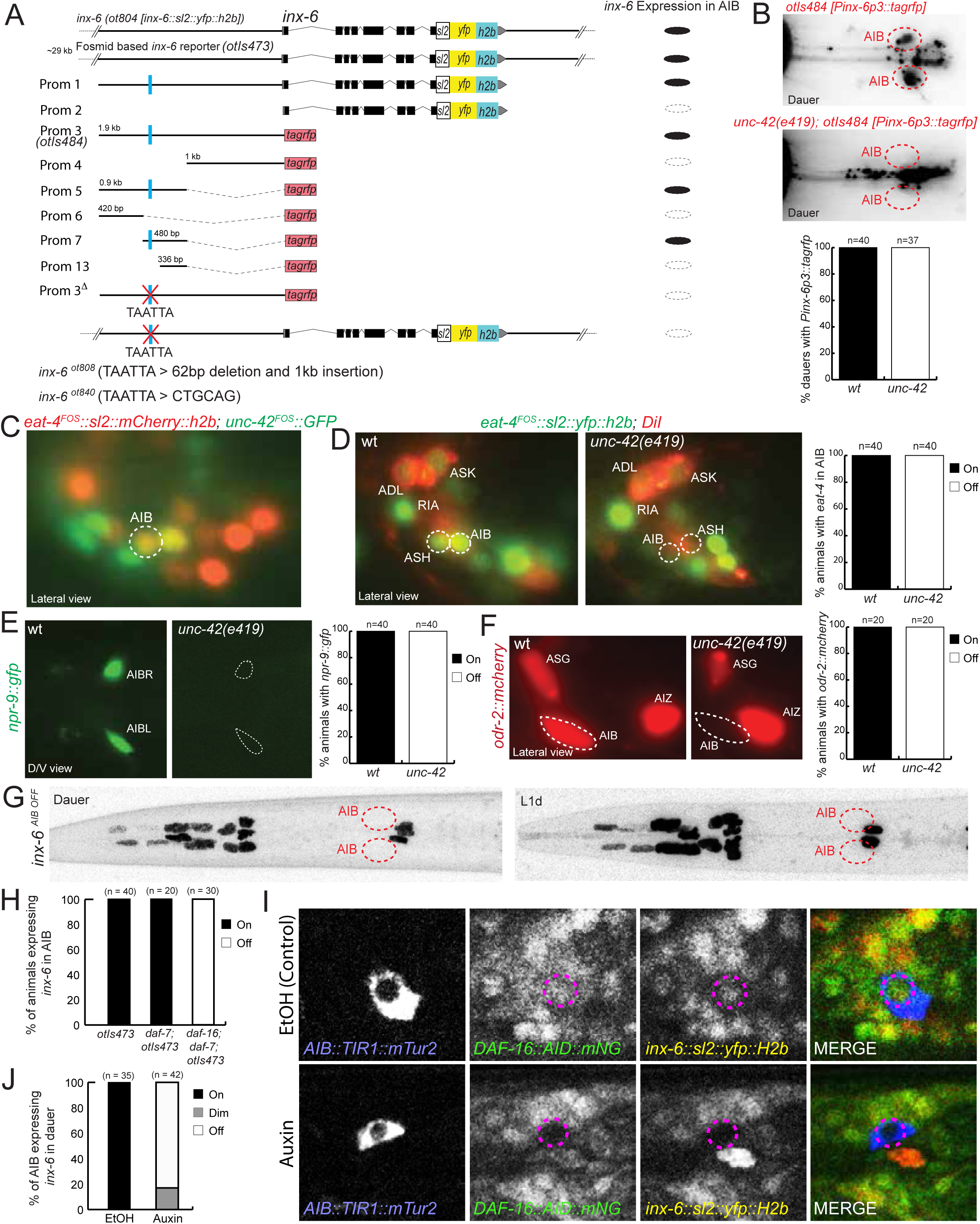
UNC-42 and DAF-16/FOXO regulate spatiotemporal expression of *inx-6* in AIB. **(A)** Analysis of 5’ cis-regulatory elements of *inx-6*. For the cis-regulatory element analysis, 2 to 3 independent transgenic reporter lines were scored (n≥25 per line). Lines commonly showed very similar expressivity and penetrance. Average results for all transgenic lines are shown. Deletion of a putative UNC-42 binding site (TAATTA) in the 5’ upstream regulatory region using CRISPR/Cas9-mediated genome editing resulted in complete loss of *inx-6* expression in AIB. Strains are listed in Table S5. **(B)** UNC-42 affects *inx-6* reporter (*otIs484[inx-6p3::tagrfp]*) expression in AIB in dauer stage. % of animals expressing *inx-6* reporter in AIB are indicated. **(C)** A fosmid-based GFP-tagged *unc-42* reporter transgene *(wgIs173[unc-42*^*fosmid*^::*gfp])* (green) is expressed in AIB interneurons in all stages. Neuronal identities were determined by expression of *otIs518[eat-4*^*fosmid*^::*sl2::mcherry::H2B]* reporter (red). **(D-F)** Expression of multiple terminal identity markers of AIB, *eat-4, npr-9 and odr-2,* were affected in *unc-42(e419)* mutant animals. **(G)** Deletion of a putative UNC-42 binding site in *inx-6(ot840)* animals, results in loss of *inx-6* expression specifically in AIB neurons in dauer and L1-diapause. **(H)** Quantification of fosmid-based *inx-6* transcriptional reporter (*otIs473[inx-6*^*fosmid*^::*sl2::yfp:: H2B])* expression in dauer. *inx-6* expression is lost in AIB in *daf-7(e1372); daf-16(mgDf50)* double mutant dauers. **(I)** AIB specific degradation of DAF-16::mNG::AID by neuron specific expression of TIR1::mTurquoise2. In auxin-treated dauer animals, DAF-16::mNG::AID protein expression as well as *inx-6* expression (as assessed by *inx-6(ot804)* reporter allele expression) are lost in AIB. In EtOH-treated control dauer animals, expression of DAF-16, as well as *inx-6* are maintained. Due to substantial overlap of mNeonGreen and YFP emission spectra these two expressions could not be separately imaged. *(inx-6(ot804); daf-2(e1370); daf-16(ot853[daf-16::linker::mNG::3xFlag::AID]; otEx7309[inx-1p::TIR1::unc-54 3’UTR::rps-27p::NeoR::unc-54 3’UTR])* **(J)** Quantification of results shown in panel I.

*unc-42* not only affects the dauer-specific *inx-6* expression in AIB but also affects other genes expressed constitutively in both dauer and non-dauer AIB neurons. Specifically, we found that *unc-42* is required for the expression of *eat-4/VGLUT,* neuropeptide receptor encoding gene *npr-9* and an olfactory signaling protein encoding gene *odr-2* (Fig. 7D-F), indicating that *unc-42* may be a terminal selector of AIB identity.

The dauer-specificity of *inx-6* induction suggests that *unc-42* alone is not sufficient to turn on *inx-6* expression but requires dauer-specific regulatory inputs. To investigate those, we turned to the insulin/IGF-1 signaling pathway, a major determinant of the dauer arrest mechanism (Fielenbach and Antebi, 2008). In non-dauer animals, the insulin/IGF-1 signaling pathway suppresses the nuclear translocation of the DAF-16/FOXO transcription factor. In dauer animals, insulin/IGF-1 signaling is downregulated leading to nuclear translocation of DAF-16, where it regulates many different target genes in different tissue types (Fielenbach and Antebi, 2008). To assess the involvement of *daf-16,* we examined *inx-6* expression in *daf-16* mutant animals that were forced into dauer by simultaneously mutating the *daf-7/TGFβ* endocrine signal pathway (Lee et al., 2001). In these double mutant dauers *inx-6* fosmid reporter expression was not induced in AIB, demonstrating that *daf-16* is required for the dauer-specific *inx-6* expression (Fig. 7H). To assess the focus of action of DAF-16, we removed DAF-16 exclusively from the AIB neuron. To this end, we inserted an Auxin-Inducible-Degron (AID)-tag (Zhang et al., 2015) together with the fluorescent gene *mNeonGreen* at the 3’end of the *daf-16* locus, using CRISPR/Cas9-mediated genome editing. This allele allowed us to follow DAF-16 localization dynamics in the nervous system and conditionally degrade DAF-16 in a spatiotemporal manner. Upon entry into the dauer stage, DAF-16::mNG showed the expected nuclear translocation. AIB-specific expression of TIR1 in conjunction with the auxin treatment successfully depleted DAF-16 only in AIB, but did not affect dauer formation (Fig. 7I). This cell-autonomous depletion of DAF-16 in AIB prevented *inx-*6 expression in dauer (Fig. 7I,J). These results suggest that the environmental cues control the insulin/IGF-1 signaling pathway leading to altered DAF-16 activity in AIB, where it cell-autonomously induces *inx-6* expression in dauer.

## DISCUSSION

While there is a general appreciation of the importance of establishing synaptic connectivity diagrams, much of past and present analysis of connectomes focuses on chemical synapses, even though the importance of electrical synapses in nervous system function has been made apparent by a number of genetic loss of function studies (Abrams and Scherer, 2012; Hall, 2017; Hasegawa and Turnbull, 2014; Marder et al., 2017; Song et al., 2016; White and Paul, 1999). Through the establishment of a nervous system wide map of innexin expression, as well its dynamic modulation under specific environmental conditions; we have laid here the groundwork to understand a number of distinct features of the electrical connectome. First, it is generally thought that the formation of electrical synapses is encoded by the neuron-type-specific expression of matching homo- or heteromeric hemichannels, which recognize each other in *trans* to assemble into a functional electrical synapse. Second, the specific composition of an electrical synapse is thought to endow an electrical synapse with specific conductive properties (Baker and Macagno, 2017; Goodenough and Paul, 2009; Mese et al., 2007; Miller and Pereda, 2017; Sohl et al., 2005). For example, the widespread electrical connectivity of neurons in the *C. elegans* nervous system clearly indicates that information flow through many of the electrical synapses must be directional and such directionality is likely encoded by the specific molecular composition of individual synapses.

As a prerequisite to address these two issues, we identified the complex innexin expression code of the *C. elegans* nervous system. 14 out of 25 innexin genes in the *C.* elegans genome are expressed in complex combinatorial codes in the nervous system under distinct environmental conditions. Some innexins are expressed very broadly in the nervous system, some much more restrictively. Alternate splice isoforms of a single innexin can be expressed in distinct neuronal classes. Furthermore, the number of distinct innexin genes present in a particular neuronal class also varied greatly. Whereas ASK and RIM neuron classes expresses the maximum of 11 innexins, the I6 neuron class expresses only one innexin. We hypothesize that this distinct neuron-specific combinatorial innexin expression code may be a key underlying determinant for the establishment of the electrical connectome and may further provide neurons with the ability to form electrical synapses with potentially synapse-specific conductance, which depends on the specific innexin composition.

We have shown that the innexin expression code of the nervous system is not static but rather subjected to widespread, substantial alteration when developing *C. elegans* larvae encounter harsh environmental conditions and enter the dauer diapause stage. In the dauer nervous system, the expression of 12 out of 14 innexins that are expressed in the nervous system in non-dauer stages is altered in 86 out of 118 neuron classes. Additionally one non-neuronal innexin gene, *inx-6,* is also switched on in the dauer nervous system. Often the expression of a single innexin is turned on in some neuron classes while simultaneously turned off in others. Expressions of distinct splice isoforms of a single innexin are also independently altered in distinct neuronal classes. Some neuron classes simultaneously upregulate one innexin and downregulate another to change their innexin expression code. Changing the expression of innexins in a given neuron class could have two different consequences. It may lead to an alteration (gain and/or loss) of synaptic partners and/or it may lead to changes in the composition of existing synaptic contacts. In either case, these changes are predicted to have an impact on information flow in the nervous system. In one of the cases that we described here in more detail, the upregulation of *inx-6* in AIB, results in the generation of novel electrical synaptic connections. It will require the electron micrographic reconstruction of the entire nervous system of dauer stage animals to assess the extent of altered synaptic partner choice. Our expression map of innexins provides the foundation to understand the molecular mechanisms of structural rewiring observed by such anatomical analysis.

To provide functional correlates to changes in the electrical connectome, we analyzed locomotory behavior of dauers and discovered specific patterns of alteration compared to developmental stage matched non-dauer animals. Notably, although we obtained dauers under prolonged starvation condition (accompanied by high population density and temperature), some aspects of dauer behavior were more similar to satiated non-dauer animals than to short-term starved non-dauer animals, suggesting long term adaptions to starvation. Several changes in locomotory behavior are, however, clearly dauer-specific and fall in line with the previously reported behavioral alteration of dauer stage animals, such as dauer specific increased quiescence, dispersal type nictation behavior and altered chemotactic preference (Cassada and Russell, 1975; Gaglia and Kenyon, 2009; Lee et al., 2012). We have linked here one of these behavioral alterations to the generation of a novel electrical synapse, generated between the BAG sensory neuron and the AIB interneuron specifically in dauer animals. We also found that downregulation of a widely expressed innexin, *unc-7*, in one specific interneuron pair is required for some of the dauer-specific changes in locomotory behaviors.

Electrical synapse assembly and function is known to be regulated by a number of distinct mechanisms (O’Brien, 2014; Thevenin et al., 2013) and our description of gene expression changes of innexins only describes the first layer of regulation of the electrical connectome. Our characterization of such transcriptional dynamics offers a unique opportunity to study the plasticity of gene expression in mature post-mitotic neurons. We have provided here some mechanistic insights into how the gene expression plasticity of innexins is controlled. For all neuronal classes studied in *C. elegans,* expression and maintenance of the neuron type specific terminal fate makers are governed by continuously expressed terminal selector type transcription factors. We have described a paired-type homeodomain transcription factor, UNC-42, controls several invariant features of AIB identity and found the neuronal specificity of the dauer specific *inx-6* induction in AIB also depends on UNC-42. The condition-specific (i.e. dauer-specific) aspect of innexin expression could not be solely explained by the constitutive activity of these constitutively active terminal identity specifiers. We rather found this dynamic aspect of *inx-6* expression in dauer to be controlled by the cell-autonomous inactivation of insulin/IGF-1 signaling and resultant activation of DAF-16/FOXO. This intersectional gene regulatory strategy may constitute a generalizable mechanism for how the environment can control neuronal plasticity with cellular specificity.

## EXPERIMENTAL PROCEDURES

### *C. elegans* strains and transgenes

Wild-type strain used is *C. elegans* variety Bristol, strain N2. Worms were grown on nematode growth media (NGM) plates seeded with bacteria (OP50) as a food source at 20°C, unless otherwise mentioned. Worms were maintained according to standard protocol.

A list of strains and transgenes used in this study is listed in Supplementary Table S5.

### Cloning and constructs

#### Fosmid reporter clones

The sl2-based transcriptional innexin fosmid reporters was generated by recombining an SL2 trans-splicing sequence followed by a fluorescent reporter cassette containing the *yfp*-gene sequence tagged with an NLS and a histone (H2B) at the 3′ end of the respective gene locus just after the stop codon, using fosmid recombineering (Tursun et al., 2009).

#### *inx-19* fosmid extension

Fosmid clone WRM0632bE10 in pCC1FOS was digested with ApaLI and self ligated to get a subclone containing 30kb genomic fragment (*inx-19* locus, 2.3kb sequence 5′ and 8kb sequence 3′ to the *inx-19* locus). This subclone was linearized with BamHI digestion (−2320 from *inx-19* ATG) and a 9944bp genomic sequence was inserted in using Gibson assembly (NEB, Cat. # E2621S). This final fosmid clone contained *inx-19* locus, 12.3kb sequence 5′ and 8kb sequence 3′ to the *inx-19* locus.

For bi-cistronic *che-7* reporter fosmid a *tagrfp::Sl2::NLS::yfp::H2B* cassette was recombined right before the stop codon of the locus, using fosmid recombineering (Tursun et al., 2009).

The *inx-3, inx-9* and *inx-17* translational fosmid reporter constructs were kindly provided by the TransgeneOme project (Sarov et al., 2012). *gfp* was recombined right before the stop codon of the locus. A detailed list of fosmids generated is provided in the Supplementary Experimental procedure (Table S5).

#### Genome editing

Fluorescent knock-in alleles of *inx-2 and inx-6* or mutant allele of *inx-6* was generated using CRISPR/Cas9-triggered homologous recombination based on co-CRISPR method (Kim et al., 2014).

Generation of *daf-16(ot853[daf-16[daf-16::mNG::3XFLAG::AID]* allele: CRISPR/Cas9-mediated genome editing using a self-excising cassette (SEC) as previously described (Dickinson et al., 2015). We modified the pDD268 plasmid (developed in (Dickinson et al., 2015)), which contains the mNeonGreen (mNG) fluorescent reporter cassette, to include the Auxin Inducible Degron (AID)(Zhang et al., 2015) after the 3xFLAG tag. We inserted this mNG::3xFLAG::AID tag right before the stop codon of the *daf-16* gene locus to produce mNG::AID fused DAF-16 protein. This tagging captures all DAF-16 isoforms, since they have a common C-terminus.

To generate *npr-9p::tagrfp, a 1.7kb cis-*regulatory region (−1689 to 0) was fused with *tagrfp::unc-54 3′ UTR* sequence using PCR fusion.

To generate *ets-5p::tagrfp, a 3.3kb cis-*regulatory region (−3276 to 0) was fused with *tagrfp::unc-54 3′ UTR* sequence using PCR fusion.

All reporter transgenes for *cis-*regulatory analysis of *inx-6* locus were generated using a PCR fusion approach (Hobert, 2002) using *tagrfp* coding sequence. Coordinates with respect to ATG were indicated in Fig. 7A.

All reporter transgenes for *cis-*regulatory analysis of *che-7* locus were generated by amplifying indicated regions from the bi-cistronic *che-7*^*fosmid*^::*tagrfp::Sl2::NLS::yfp::H2B* fosmid clone.

For neuron specific expression of *che-7::tagrfp::Sl2::NLS::yfp::H2B,* total cDNA was synthesized using oligo(dT)_20_ primer and SuperScript III First-Strand Synthesis System, according to the Invitrogen protocol. *che-7* cDNA was amplified using primers: 5′ ATGCCAGAAAACAAACTTCAATTGG 3′ and 5′ CAAATCTAGAAGAGAACTGGC 3′ and cloned into pMiniT 2.0 vector (NEB). *tagrfp::Sl2::NLS::yfp::H2B* cassette was amplified from the bi-cistronic *che-7*^*fosmid*^::*tagrfp::Sl2::NLS::yfp::H2B* fosmid clone and introduced in the pMiniT-*che-7*^cDNA^ clone right before the *che-7* stop codon to get the pMiniT-*che-7*^*cDNA*^::*tagrfp::SL2::NLS::yfp::H2B* clone using Gibson assembly. 3311bp *flp-17p* [BAG specific], 897bp *gcy-8p* [AFD specific] and 1380bp *che-1pA* [ASE specific] was introduced right before the *che-7* ATG using Gibson assembly. *Cell specific prom::che-7*^*cDNA*^::*tagrfp::SL2::NLS::yfp::H2B::che-7 3′ UTR* was amplified from the final clone and injected to get transgenic animals (see Table S5).

To generate *flp-18p::unc-7::SL2::NLS::yfp::H2B, unc-7::SL2::NLS::yfp::H2B unc-7 3′ UTR* cassette was amplified from the *unc-7::SL2::NLS::yfp::H2B* fosmid clone using upstream primer *5′* TAACACGAACCCGGGATGCTCGGCTCCTCCAGCAA 3′ (this also introduces a SmaI cut site right before the *unc-7* ATG) and downstream primer 5′ CCTCAAATTGAGCCCATCAG 3′ and sub-cloned into pTOPO XL vector (Invitrogen). This construct was linearized using SmaI digestion and 3117bp *flp-18p* sequence was introduced before the *unc-7* coding sequence using Gibson assembly.

### Microscopy

Worms were anesthetized using 100mM of sodium azide and mounted on 5% agarose on glass slides. Images were recorded using either Zeiss 880 confocal laser-scanning microscope or Zeiss Axio Imager Z2 wide field fluorescent microscope. Images were analyzed by scanning the full Z-stack using Zeiss Zen software. Maximum intensity projections constructed using NIH Fiji software of representative images were shown (Schneider et al., 2012). Figures were prepared using Adobe Photoshop CS6 and Adobe Illustrator CS6. Separate channels were usually adjusted independently using Levels and Curves in Adobe Photoshop.

### Neuron identification

Cell identification for reporter expression was done by Nomarski optics and crossing with neuronal landmark reporter strains, primarily fosmid reporters of *eat-4 (otIs518), cho-1 (otIs544)* and *unc-47 (otIs564),* and promoter-fusion reporter of *rab-3 (otIs355)* (Gendrel et al., 2016; Pereira et al., 2015; Serrano-Saiz et al., 2013; Stefanakis et al., 2015). Expression in a subset of sensory neurons was identified additionally by dye filling with DiI (Thermo Fisher Scientific).

### *C. elegans* electrical synapse connectome

The electrical synapse network diagram was drawn using open source software, Cytoscape (www.cytoscape.org) based on synaptic connectivity data obtained from serial section reconstruction of electron micrographs (TEM) collected in MRC/LMB and rescored in Emmons and Hall lab (Jarrell et al., 2012; White et al., 1986).

### single molecule fluorescence in situ hybridization (smFISH)

smFISH analysis was performed as previously described (Ji and van Oudenaarden, 2012). Samples were incubated over night at 37°C during the hybridization step. The *inx-6* probes were designed by using the Stellaris RNA FISH probe designer. Purified probes conjugated to Quasar 670 dye were obtained from Biosearch Technologies.

### Automated worm tracking

Automated single worm tracking was performed using Wormtracker 2.0 system at room temperature (∼ 22°C) except for temperature sensitive *inx-6(rr5)* strain, which were performed at 25°C (Yemini et al., 2013). Dauer animals, which tend to show extensive pausing, were recorded for 10 min to ensure sufficient sampling of locomotion related behavioral features. Non-dauer animals were recorded for 5 min, except when compared to dauer animals (as in Fig. 1) were recorded for 10 min. To avoid potential bias towards pausing, worms that were actively moving were selected for tracking. To avoid potential variability arising due to room conditions, all strains that were compared in a single experiment were recorded simultaneously in identical room condition, along with N2 wild-type. Strains that were recorded simultaneously with temperature sensitive *inx-6(rr5)* strain, were grown at 25°C for >24 hr and recorded at 25°C. Recording was randomized across multiple trackers. Analysis of the tracking videos was performed as previously described. Dauer and starved non-dauer animals were placed on uncoated NGM plates before recording. Fed non-dauer animals were recorded on NGM plates seeded uniformly with diluted OP50 bacterial culture to avoid potentially biased locomotion at the edge of the bacterial lawn. For the comparison of dauer locomotion with fed and starved L3 locomotion, 702 behavioral features were measured. After correction for multiple testing (Bonferroni correction), features shown in Fig. 5 and Table S4 immerged as the once with most significantly different q-value among the test groups. Therefore, for the subsequent experiments we only measured these features, permitting us to use the p-value. The principle components analysis (PCA) was executed using MATLAB (Mathworks) with custom computational scripts (made available upon request).

### CO_2_ Chemotaxis assay

CO_2_ chemotaxis assays were performed as previously described (Hallem et al., 2011a). Assays were performed on standard 6cm NGM plates for non-dauer animals and 9cm NGM plates for dauer animals. Scoring regions were 2cm circles on each side of the plate along the diameter, with the center of the circle 1cm away from the edge of the plate (as indicated in Fig. 6J). Pumping a mixture of 10% CO2, 10% O2 and balance N2 wild-type from an inlet on top of one scoring circle and a mixture of 10% O2 and balance N2 wild-type from another inlet on top of the other scoring circle generated CO2 gradient. Gas mixtures were pushed using a syringe pump at 1.5ml/min for non-dauer assays and 0.5ml/min for dauer assays. ∼50 young adults or ∼100-150 dauers (selected by 1% SDS treatment) were placed at the center of the assay plates. After 30 min, the number of animals inside the air and CO_2_ circles was counted. The chemotaxis index (C.I.) was calculated as:

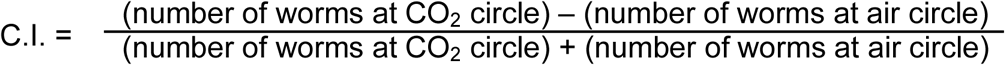

### Odortaxis assay

Dauer odortaxis assays were performed as previously described (Hallem et al., 2011a). Assays were performed on standard 9cm NGM plates. Scoring regions were 2cm circles on each side of the plate along the diameter, with the center of the circle 1cm away from the edge of the plate. Test circle contained 5μl of odorant at center of it, while control circle contained 5μl of ethanol. 1μl of 1M sodium azide was added to each circle as anesthetic. ∼100-150 dauers were placed at the center of the assay plates and allowed to chemotax for 90min.

The chemotaxis index (C.I.) was calculated as:

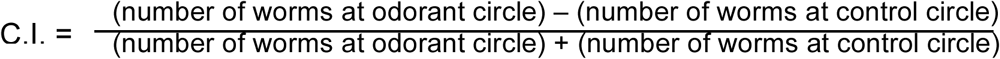

### Cell ablation

Cell specific ablation was performed using MicroPoint Laser System Basic Unit (N2 pulsed laser (dye pump), ANDOR Technology) attached to a Zeiss Axio Imager Z2 microscope (Objective EC Plan-Neofluar 63X/1.30 Oil). This laser delivers 120 μJoules of 337nm energy with a 3nsec pulse length. Ablations were done as previously described (Fang-Yen et al., 2012), with pulse repetition rates ∼15 Hz. *daf-7(e1372); npr-9::tagrfp; inx-6(ot805)* stain was grown at 25°C to obtain dauer. AIB identified with TAGRFP reporter and ablated at the dauer stage. Treated animals were allowed to recover at 25°C on unseeded NGM plates for 48 hours before analysis. Control animals were treated in the same manner, but not subjected laser exposure.

### Auxin inducible degradation

The AID system was employed as previously described (Zhang et al., 2015). The conditional *daf-16* allele *daf-16(ot853[daf-16::mNG::AID)* was crossed with *daf-2(e1370)*, neuron specific TIR1-expressing transgenic lines and *inx-6(ot804[inx-6::sl2::yfp::h2b]* reporter, to generate the experimental strains. Animals were grown (from embryo onwards) on NGM plates supplemented with OP50 and 4mM auxin in EtOH (indole-3 acetic acid, IAA, Alfa Aesar, Cat. # A10556) at 25°C to degrade DAF-16 in respective neurons and to induce dauer formation. As control, plates were supplemented with the solvent EtOH instead of auxin.

## ACKNOWLEDGMENTS

We thank Niels Ringstad and Jung-Hwan Choi for introducing us to the CO2 assay, Esther Serrano-Saiz and Narmin Tahirova for their help with the *unc-42* mutant analysis, Adriane Otopalic for helping with the PCA, Steven Cook for helping with the connectome, Chi Chen for the *C. elegans* injections and members of the Hobert lab for comments on this manuscript. This work was supported by the NIH (R21NS106909) and the Howard Hughes Medical Institute.

## Supplementary Figure Legends

**Fig. S1:**
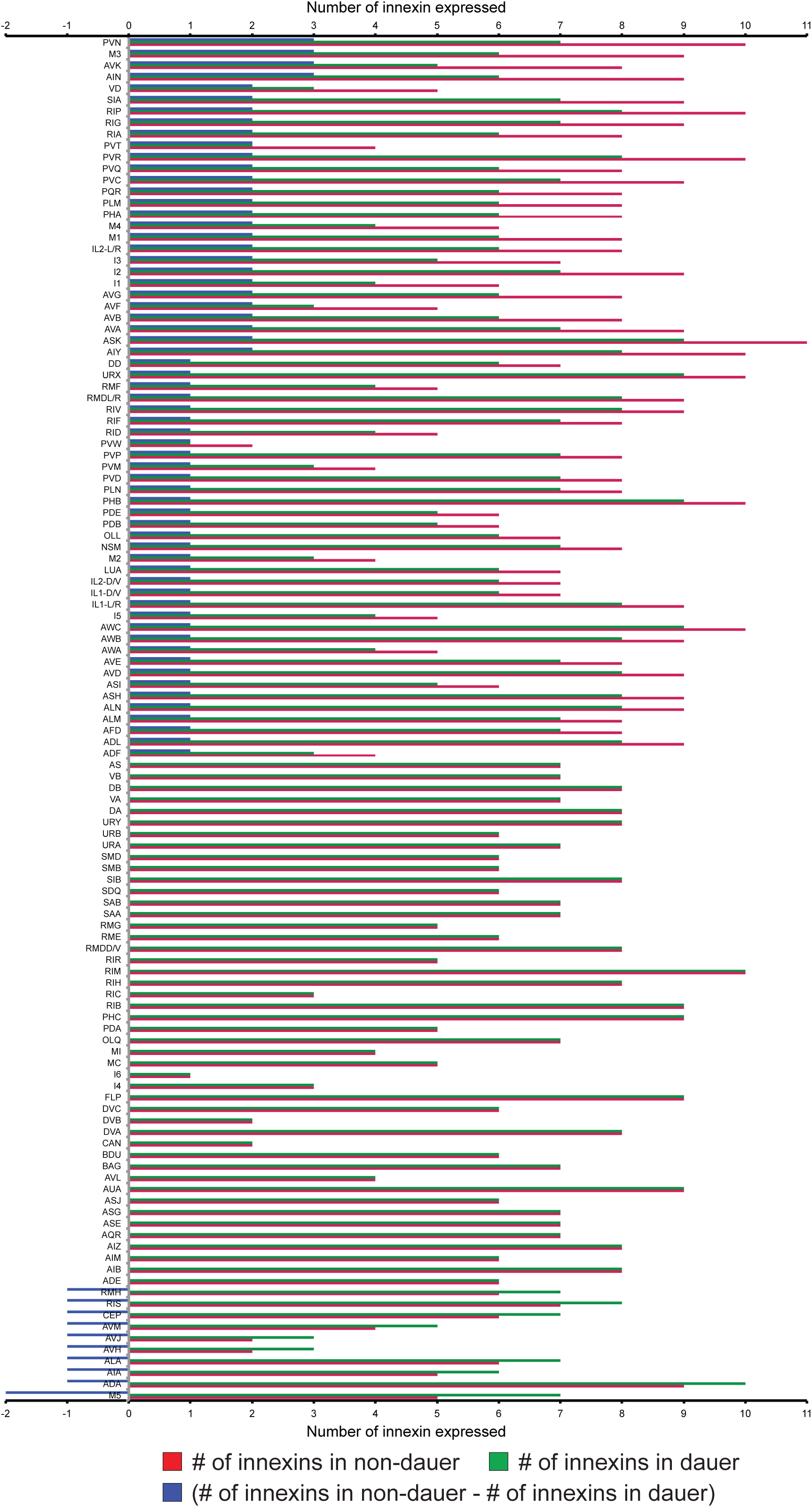
innexin expression difference in dauer and non-dauer nervous system. Graphical representation of innexin expression differences in each neuron classes. HSN and VC neurons, which mature only at L4 stage and showed no innexin expression in dauer, were not included in this graph.

**Fig. S2:**
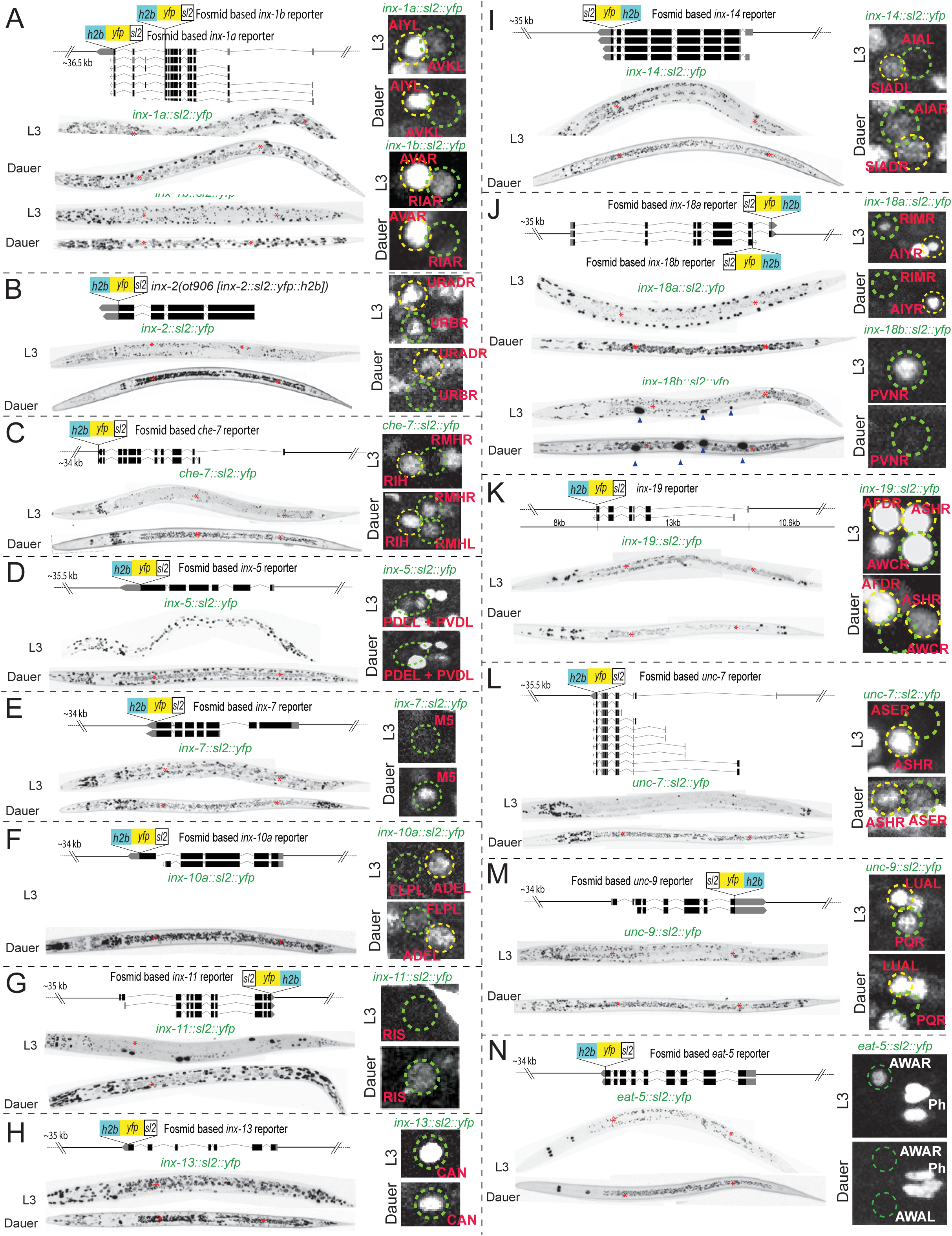
Expression of neuronally expressed innexins change in dauer stage. Schematics of fosmid-based transcriptional reporters of neuronally expressed innexin genes are shown. For *inx-2,* schematic of an sl2-based transcriptional reporter allele, *inx-6(ot906)* generated by CRISPR/Cas9-mediated genome editing, is shown. Splice isoform specific fosmid reporters for *inx-1, inx-10* and *inx-18* are generated. Whole animal expressions for each innexin gene in L3 and dauer stages are shown. Red asterisks mark gut auto-fluorescence. Examples show selective up or down-regulation of particular innexin gene expression in specific neuronal classes, as assessed by the indicated reporters. *inx-13*, which does not show altered expression in dauer, continues to be expressed in CAN neurons in dauer. Strains are listed in Table S5.

**Fig. S3:**
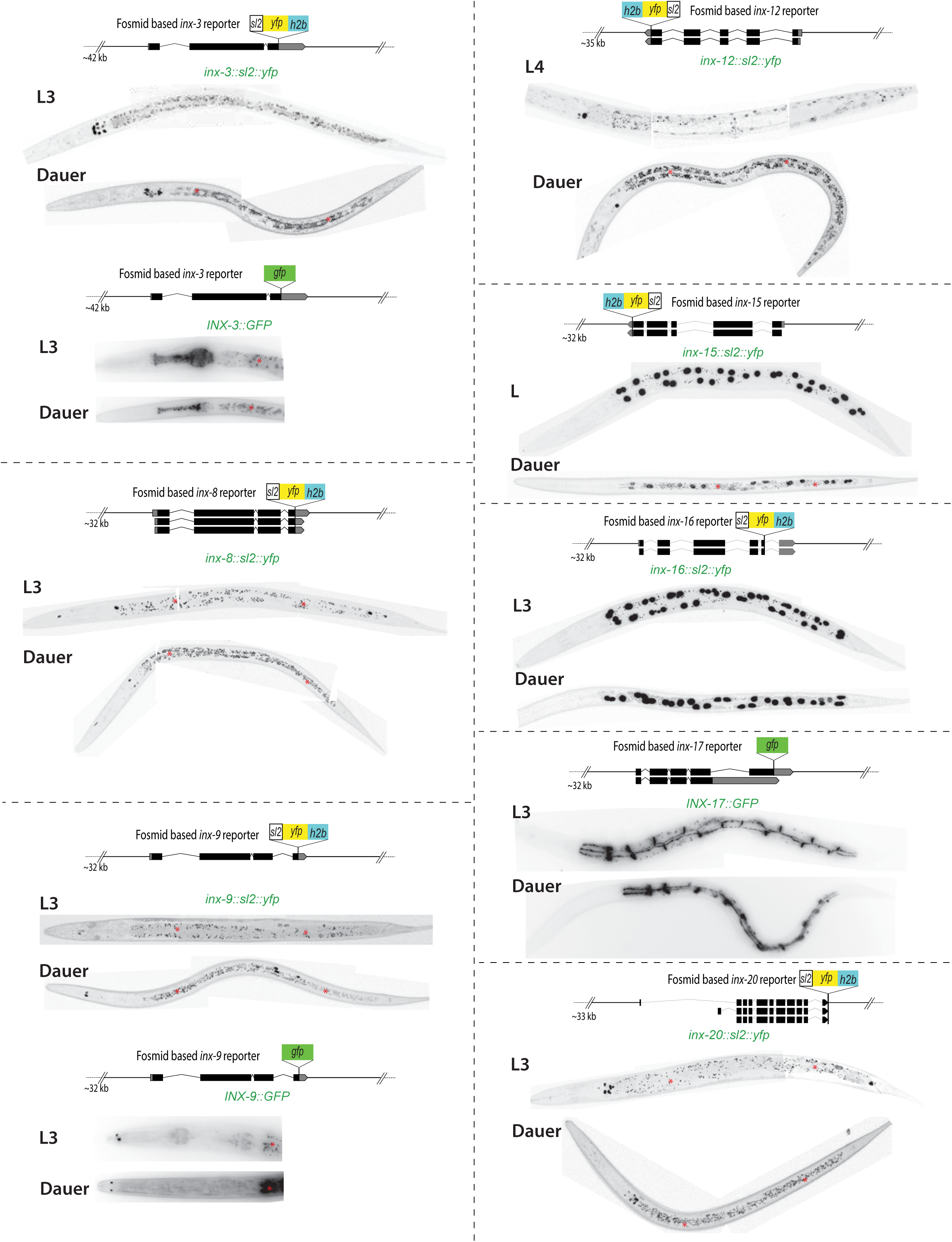
Expression of non-neuronal innexin genes. Schematics of fosmid-based transcriptional reporters of non-neuronally expressed innexin genes are shown. For, *inx-3, inx-9* and *inx-17* schematics of fosmid-based GFP-tagged translational reporters are shown. Whole animal expressions for each innexin gene in L3 and dauer stages are shown. For, *inx-3* and *inx-17* GFP-tagged translational reporters only the relevant head expression is shown. Red asterisks mark gut auto-fluorescence. Strains are listed in Table S5.

**Fig. S4:**
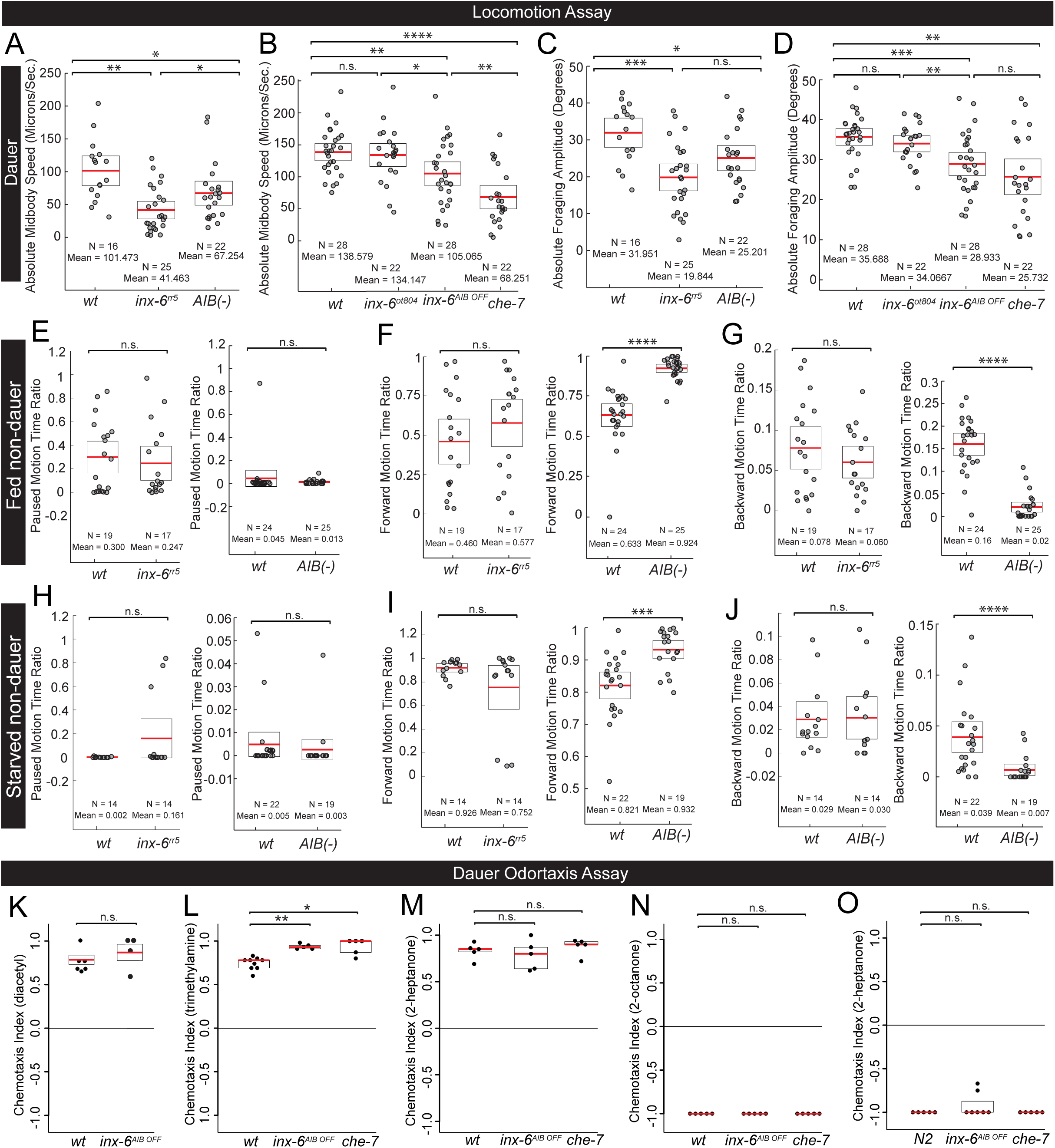
*inx-6* and *che-7* affect locomotory behavior specifically in dauer stage. **(A-J)** Worm locomotion assay using Wormtracker 2.0 system (see Methods for detailed description). Each circle represents the experimental mean for a single animal for the corresponding behavioral assay. Red lines indicate the mean of means and black open rectangles indicate S.E.M. for each assay. Wilcoxon rank-sum tests p-values for each comparison (n.s. = non significant, *p<0.05, **p<0.01, ***p<0.001, ****p<0.0001). **(A)** Dauers with a temperature sensitive allele of *inx-6, inx-6(rr5)* and AIB-ablated dauers (*peIs578[npr-9p::casp1, npr-9p::venus, unc-122p::mCherry])* shows significant decrease in speed (measured as midbody speed). **(B)** Dauers with an *inx-6* transcriptional reporter allele, *inx-6*^*ot804*^, shows no significant effect in speed. Dauers that specifically lack *inx-6* expression in AIB, *inx-6*^*AIB*^ ^*OFF*^ and *che-7(ok2373)* dauers shows significant decrease in speed (measured as midbody speed). **(C)** Dauers with a temperature sensitive allele of *inx-6, inx-6(rr5)* and AIB-ablated dauers (*peIs578[npr-9p::casp1, npr-9p::venus, unc-122p::mCherry])* shows significant decrease in foraging amplitude. **(D)** Dauers with an *inx-6* transcriptional reporter allele, *inx-6*^*ot804*^, shows no significant effect in foraging amplitude. Dauers that specifically lack *inx-6* expression in AIB, *inx-6*^*AIB*^*OFF* and *che-7(ok2373)* dauers shows significant decrease in foraging amplitude. **(E-J)** In fed-L3 and starved-L3 stage animals, *inx-6(rr5)* animals shows no significant affect on paused, forward and backward motion time ratio, compared to N2 wild-type animals. AIB-ablated fed- and starved-L3 animals (*peIs578[npr-9p::casp1, npr-9p::venus, unc-122p::mCherry])* shows no difference in pausing, but strong reduction in backward locomotion and subsequent increase in forward locomotion. **(K-O)** Odortaxis assay of N2 wild-type dauer animals. Each circle represents chemotaxis index calculated from each assay. Red line indicates the mean and black rectangles indicate S.E.M. Wilcoxon rank-sum tests p-values for each comparison (n.s. = non significant, *p<0.05, **p<0.01, ***p<0.001, ****p<0.0001) (see Methods for detailed description of behavioral assays). ***(K)*** *inx-6*^*AIB*^ ^*OFF*^ dauers does not affect attraction towards diacetyl (2,3-butanedione). ***(L)*** *inx-6*^*AIB*^ ^*OFF*^ and *che-7(ok2373)* dauers show significant increase trimethylamine attraction. ***(M)*** *inx-6*^*AIB*^ ^*OFF*^ and *che-7(ok2373)* dauers does not affect 2-butanone attraction. ***(N)*** *inx-6*^*AIB*^ ^*OFF*^ and *che-7(ok2373)* dauers does not affect 2-octanone repulsion. ***(O)*** *inx-6*^*AIB*^ ^*OFF*^ and *che-7(ok2373)* dauers does not affect 2-heptanone repulsion.

**Supplementary Table 1:** Innexin expression in non-neuronal tissues

| **innexin genes** | **Reporters used** | **Expression in non-neuronal cell types** |
| --- | --- | --- |
| *inx-1a* | Transcriptional Fosmid reporter | Muscles (pharynx, body wall, vulval), hypodermis |
| *inx-1b* | Transcriptional Fosmid reporter | Muscles (pharynx, body wall, vulval), hypodermis |
| *inx-2* | Transcriptional CRISPR reporter | Pharyngeal muscles (terminal bulb) |
| *inx-3* | Transcriptional and translational Fosmid reporter | Pharyngeal muscles (isthmus & terminal bulb), uterus |
| *che-7* | Transcriptional Fosmid reporter | Few pharyngeal muscles |
| *inx-5* | Transcriptional Fosmid reporter | Broadly expressed in glial cells |
| *inx-6* | Transcriptional and translational Fosmid and CRISPR reporter | Pharyngeal muscles (corpus & MC) |
| *inx-7* | Transcriptional Fosmid reporter | Hypodermis, gonad and pharyngeal muscles |
| *inx-8* | Transcriptional Fosmid reporter | Gonadal sheath cells, DTC, hypodermis |
| *inx-9* | Transcriptional and translational Fosmid reporter | 3 corpus pharyngeal muscles (Translational fosmid shows 2 puncta in the nose), gonadal sheath cells |
| *inx-10a* | Transcriptional Fosmid reporter | Pharyngeal muscles, pharyngeal-intestinal valve, hypodermis, spermatheca, gonadal sheath cells, DTC, anal sphincter muscle, anal derepressor muscle |
| *inx-11* | Transcriptional Fosmid reporter | Pharyngeal muscles, vulval muscle, oocyte, vulval precursor cells, glia |
| *inx-12* | Transcriptional Fosmid reporter | Hypodermis, excretory cell, glia, gonadal muscles |
| *inx-13* | Transcriptional Fosmid reporter | Hypodermis, excretory cell, glia, oocyte, vulval cells (muscles, uterine cells), anal muscles |
| *inx-14* | Transcriptional Fosmid reporter | oocyte (Starich, 2014) |
| *inx-15* | Transcriptional Fosmid reporter | gut only |
| *inx-16* | Transcriptional Fosmid reporter | gut only |
| *inx-17* | Translational Fosmid reporter | gut only |
| *inx-18a* | Transcriptional Fosmid reporter | Hypodermis |
| *inx-18b* | Transcriptional Fosmid reporter | Hypodermis |
| *inx-19* | Transcriptional Fosmid reporter | Stomato-intestinal muscles |
| *inx-20* | Transcriptional Fosmid reporter | Pharyngeal muscles (terminal bulb), pharyngeal-intestinal valve, anal sphincter muscle |
| *inx-21* | Previous study (Starich, 2014) | Exclusively in gonad by antibody staining (Starich, 2014) |
| *inx-22* | Previous study (Starich, 2014) | Exclusively in gonad by antibody staining (Starich, 2014) |
| *eat-5* | Transcriptional Fosmid reporter | Pharyngeal muscles (corpus & MC), vulval epithelium |
| *unc-7* | Transcriptional Fosmid reporter | Pharyngeal muscles (corpus), uterus |
| *unc-9* | Transcriptional Fosmid reporter | Muscles (pharyngeal and bodywall), hypodermis |

**Supplementary Table 2:** Developmental changes in innexin expression

| innexin | Developmental changes in expression |
| --- | --- |
| *inx-1a* | HSN, VC (expressed L4 onwards) |
| *inx-1b* | VC (expressed L4 onwards) |
| *inx-2* | AFD, AIN (expressed until L4) |
| *inx-5* | PDE, PVD (expressed until L3) |
| *inx-7* | VC (expressed L4 onwards) |
| *inx-10a* | HSN, VC (expressed L4 onwards) |
| *inx-11* | PQR, URY (expressed until L2) |
| *inx-14* | ASJ, AWC (expressed until L4), RIA (expressed until L3),<br>RMH (expressed L4 onwards) |
| *inx-18a* | AIB, ALA (expressed until L3), PHB (expressed until L4) |
| *inx-19* | AVA (expressed in adults), RMDL/R, AIY (lost in old adults) |
| *unc-7* | HSN (expressed L4 onwards) |
| *unc-9* | HSN, VC (expressed L4 onwards) |

**Supplementary Table 3:** List of behavioral features used in PCA

|  |  |  |
| --- | --- | --- |
| **POSTURE** | Inter Forward Time | Omega Turns Time Ratio |
| Absolute Head Bend Mean | Paused Motion Frequency | Absolute Omega Turn Time |
| Absolute Forward Head Bend Mean | Paused Motion Time Ratio | Absolute Inter Omega Time |
| Absolute Paused Head Bend Mean | Paused Motion Distance Ratio | Upsilon Turns Frequency |
| Absolute Backward Head Bend Mean | Paused Time | Upsilon Turns Time Ratio |
| Absolute Neck Bend Mean | Paused Distance | Absolute Upsilon Turn Time |
| Absolute Forward Neck Bend Mean | Inter Paused Time | Absolute Inter Upsilon Time |
| Absolute Paused Neck Bend Mean | Backward Motion Frequency | Absolute Foraging Amplitude |
| Absolute Backward Neck Bend Mean | Backward Motion Time Ratio | Absolute Forward Foraging Amplitude |
| Absolute Midbody Bend Mean | Backward Motion Distance Ratio | Absolute Paused Foraging Amplitude |
| Absolute Forward Midbody Bend Mean | Backward Time | Absolute Backward Foraging Amplitude |
| Absolute Paused Midbody Bend Mean | Backward Distance | Absolute Foraging Amplitude |
| Absolute Backward Midbody Bend Mean | Inter Backward Time | Absolute Forward Foraging Amplitude |
| Absolute Hips Bend Mean | Absolute Head Tip Speed | Absolute Paused Foraging Amplitude |
| Absolute Forward Hips Bend Mean | Absolute Forward Head Tip Speed | Absolute Backward Foraging Amplitude |
| Absolute Paused Hips Bend Mean | Absolute Paused Head Tip Speed | Absolute Foraging Speed |
| Absolute Backward Hips Bend Mean | Absolute Backward Head Tip Speed | Absolute Forward Foraging Speed |
| Absolute Tail Bend Mean | Absolute Head Speed | Absolute Paused Foraging Speed |
| Absolute Forward Tail Bend Mean | Absolute Forward Head Speed | Absolute Backward Foraging Speed |
| Absolute Paused Tail Bend Mean | Absolute Paused Head Speed | **PATH** |
| Absolute Backward Tail Bend Mean | Absolute Backward Head Speed | Path Range |
| Absolute Head Bend S.D. | Absolute Midbody Speed | Forward Path Range |
| Absolute Forward Head Bend S.D. | Absolute Forward Midbody Speed | Paused Path Range |
| Absolute Paused Head Bend S.D. | Absolute Paused Midbody Speed | Backward Path Range |
| Absolute Backward Head Bend S.D. | Absolute Backward Midbody Speed | Absolute Path Curvature |
| Absolute Neck Bend S.D. | Absolute Tail Speed | Absolute Forward Path Curvature |
| Absolute Forward Neck Bend S.D. | Absolute Forward Tail Speed | Absolute Paused Path Curvature |
| Absolute Paused Neck Bend S.D. | Absolute Paused Tail Speed | Absolute backward Path Curvature |
| Absolute Backward Neck Bend S.D. | Absolute Backward Tail Speed | **EIGEN WORM** |
| Absolute Midbody Bend S.D. | Absolute Tail Tip Speed | Absolute Eigen Projection 1 |
| Absolute <sup>Forward</sup> Midbody Bend S.D. | Absolute Forward Tail Tip Speed | Absolute Forward Eigen Projection 1 |
| Absolute Paused Midbody Bend S.D. | Absolute Paused Tail Tip Speed | Absolute Paused Eigen Projection 1 |
| Absolute Backward Midbody Bend S.D. | Absolute Backward Tail Tip Speed | Absolute Backward Eigen Projection 1 |
| Absolute Backward Tail Bend S.D. | Absolute Head Tip Motion Direction | Absolute Eigen Projection 2 |
| Max Amplitude | Absolute Forward Head Tip Motion Direction | Absolute Forward Eigen Projection 2 |
| Forward Max Amplitude | Absolute Paused Head Tip Motion Direction | Absolute Paused Eigen Projection 2 |
| Paused Max Amplitude | Absolute Backward Head Tip Motion Direction | Absolute Backward Eigen Projection 2 |
| Backward Max Amplitude | Absolute Head Motion Direction | Absolute Eigen Projection 3 |
| Amplitude Ratio | Absolute Forward Head Motion Direction | Absolute Forward Eigen Projection 3 |
| Forward Amplitude Ratio | Absolute Paused Head Motion Direction | Absolute Paused Eigen Projection 3 |
| Paused Amplitude Ratio | Absolute Backward Head Motion Direction | Absolute Backward Eigen Projection 3 |
| Backward Amplitude Ratio | Absolute Midbody Motion Direction | Absolute Eigen Projection 4 |
| Eccentricity | Absolute Forward Midbody Motion Direction | Absolute Forward Eigen Projection 4 |
| Forward Eccentricity | Absolute Paused Midbody Motion Direction | Absolute Paused Eigen Projection 4 |
| Paused Eccentricity | Absolute Backward Midbody Motion Direction | Absolute Backward Eigen Projection 4 |
| Backward Eccentricity | Absolute Tail Motion Direction | Absolute Eigen Projection 5 |
| Bend Count | Absolute Forward Tail Motion Direction | Absolute Forward Eigen Projection 5 |
| Forward Bend Count | Absolute Paused Tail Motion Direction | Absolute Paused Eigen Projection 5 |
| Paused Bend Count | Absolute Backward Tail Motion Direction | Absolute Backward Eigen Projection 5 |
| Backward Bend Count | Absolute Tail Tip Motion Direction | Absolute Eigen Projection 6 |
| Coils Frequency | Absolute Forward Tail Tip Motion Direction | Absolute Forward Eigen Projection 6 |
| Coils Time Ratio | Absolute Paused Tail Tip Motion Direction | Absolute Paused Eigen Projection 6 |
| Coil Time | Absolute Backward Tail Tip Motion Direction | Absolute Backward Eigen Projection 6 |
| Inter Coil Time | Absolute Head Crawling Amplitude |  |
| Absolute Tail-To-Head Orientation | Absolute Forward Head Crawling Amplitude |  |
| Absolute Forward Tail-To-Head Orientation | Absolute Backward Head Crawling Amplitude |  |
| Absolute Paused Tail-To-Head Orientation | Absolute Midbody Crawling Amplitude |  |
| Absolute Backward Tail-To-Head Orientation | Absolute Forward Midbody Crawling Amplitude |  |
| Absolute Head Orientation | Absolute Backward Midbody Crawling Amplitude |  |
| Absolute Forward Head Orientation | Absolute Tail Crawling Amplitude |  |
| Absolute Paused Head Orientation | Absolute Forward Tail Crawling Amplitude |  |
| Absolute Backward Head Orientation | Absolute Backward Tail Crawling Amplitude |  |
| Absolute Tail Orientation | Absolute Head Crawling Frequency |  |
| Absolute Forward Tail Orientation | Absolute Forward Head Crawling Frequency |  |
| Absolute Paused Tail Orientation | Absolute Backward Head Crawling Frequency |  |
| Absolute Backward Tail Orientation | Absolute Midbody Crawling Frequency |  |
| **MOTION** | Absolute Forward Midbody Crawling Frequency |  |
| Forward Motion Frequency | Absolute Backward Midbody Crawling Frequency |  |
| Forward Motion Time Ratio | Absolute Tail Crawling Frequency |  |
| Forward Motion Distance Ratio | Absolute Forward Tail Crawling Frequency |  |
| Forward Time | Absolute Backward Tail Crawling Frequency |  |
| Forward Distance | Omega Turns Frequency |  |

**Supplementary Table 4:**
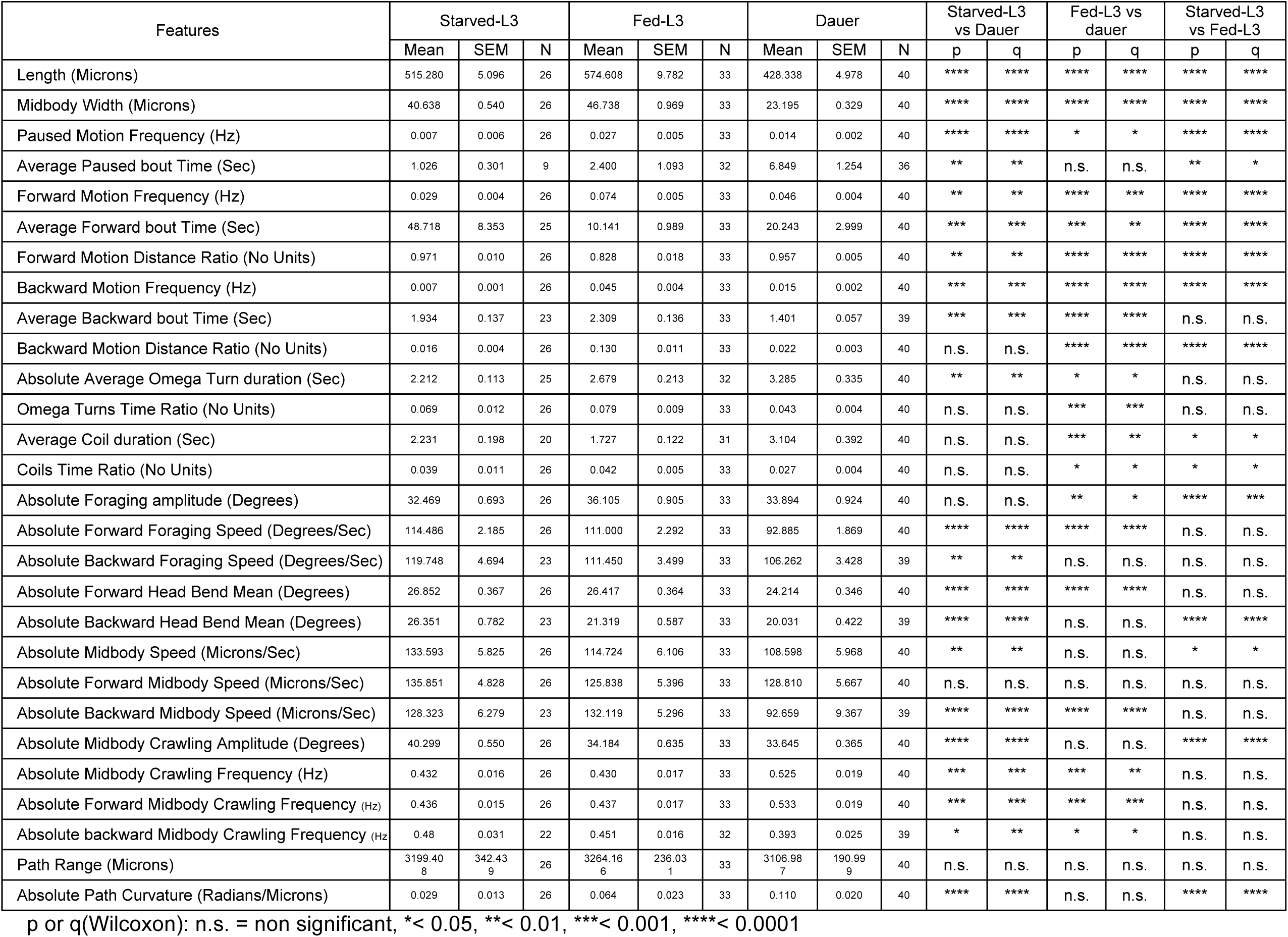
List of locomotory behavioral differences between N2 animals in dauer, fed- and starved-L3 stage/state.

**Supplementary Table 5:**
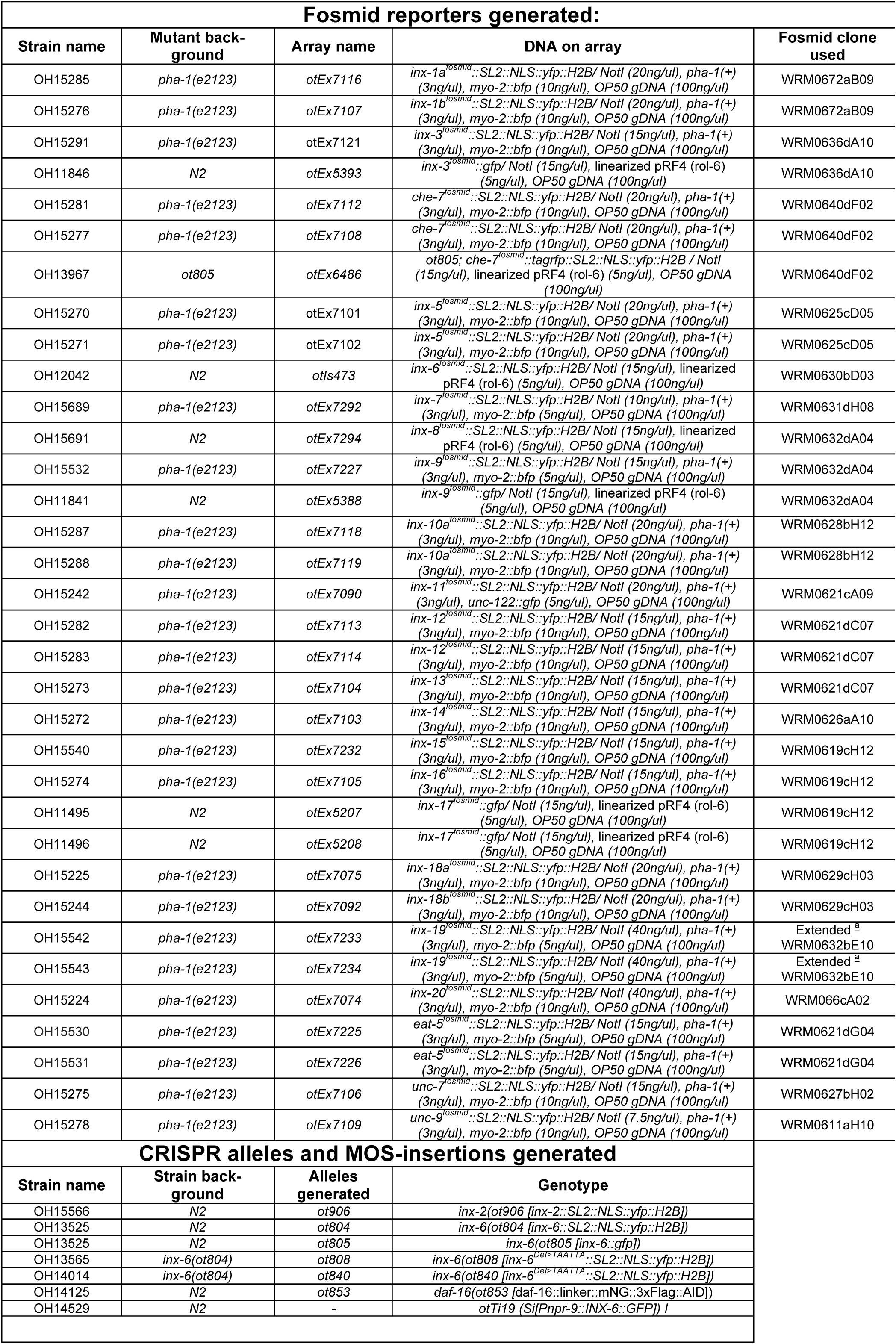
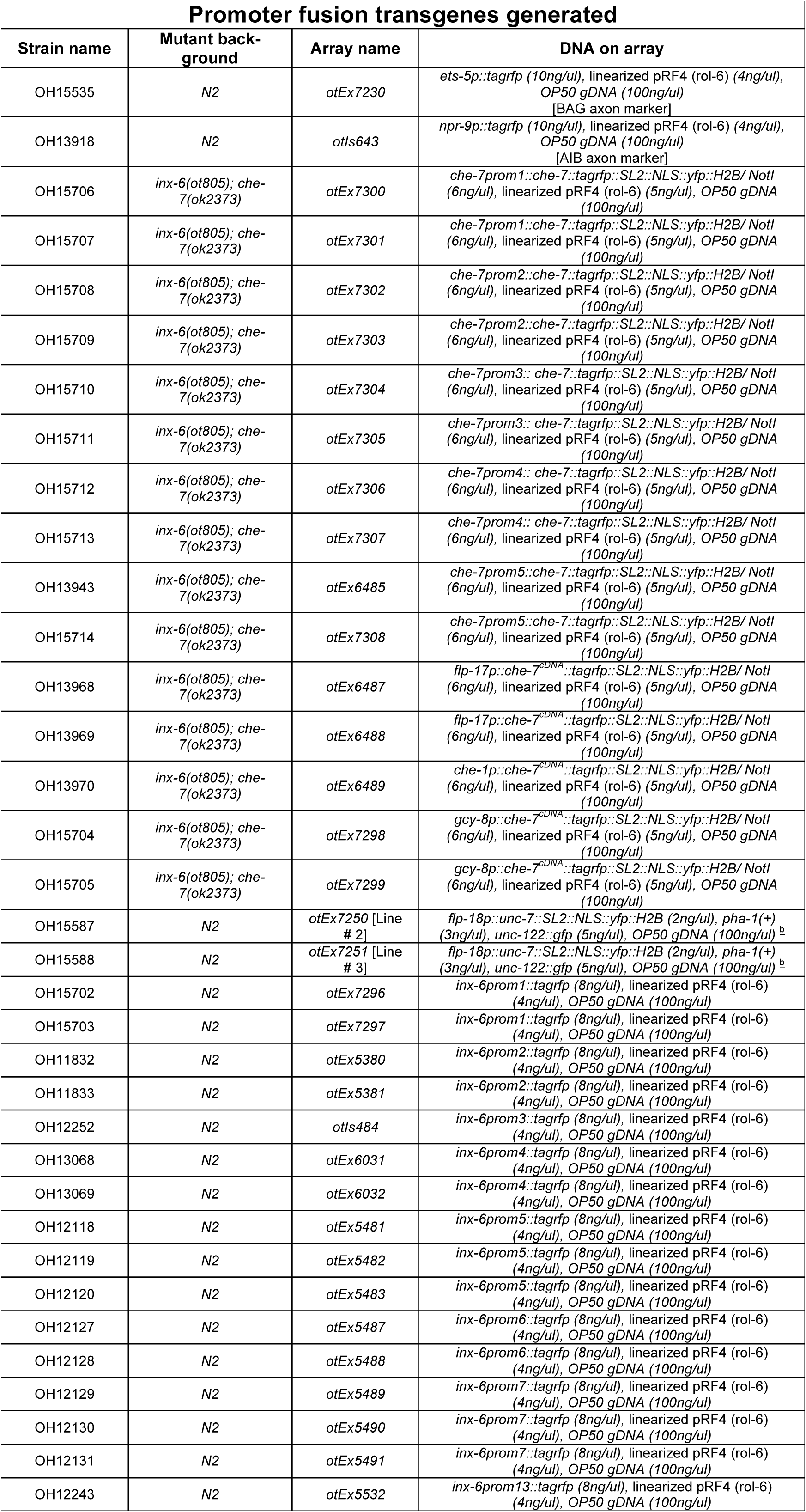
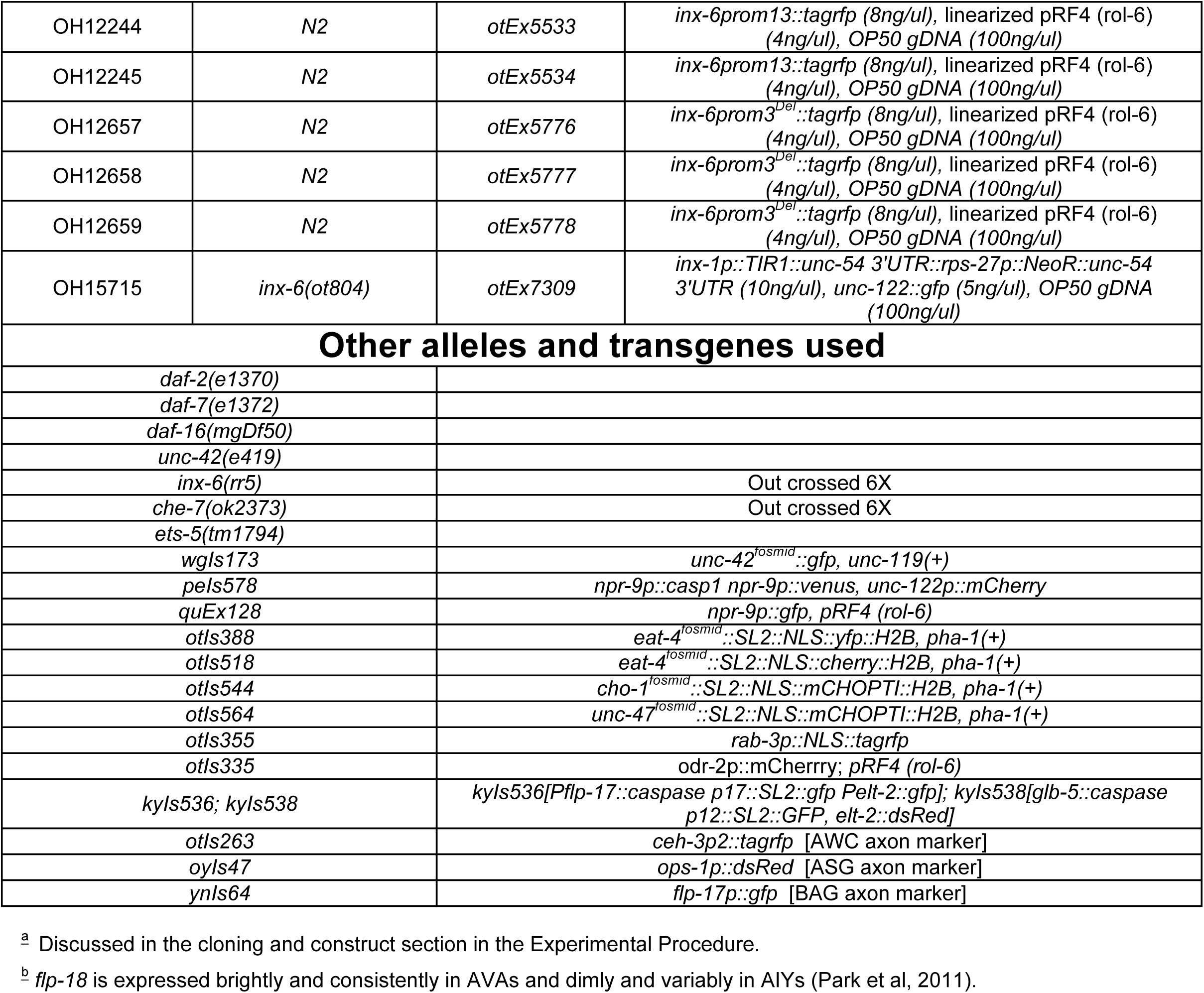
Strains used in this study

